# Systematic over-crediting in California’s forest carbon offsets program

**DOI:** 10.1101/2021.04.28.441870

**Authors:** Grayson Badgley, Jeremy Freeman, Joseph J. Hamman, Barbara Haya, Anna T. Trugman, William R.L. Anderegg, Danny Cullenward

## Abstract

Carbon offsets are widely used by individuals, corporations, and governments to mitigate their greenhouse gas emissions on the assumption that offsets reflect equivalent climate benefits achieved elsewhere. These climate-equivalence claims depend on offsets providing “additional” climate benefits beyond what would have happened, counterfactually, without the offsets project. Here, we evaluate the design of California’s prominent forest carbon offsets program and demonstrate that its climate-equivalence claims fall far short on the basis of directly observable evidence. By design, California’s program awards large volumes of offset credits to forest projects with carbon stocks that exceed regional averages. This paradigm allows for adverse selection, which could occur if project developers preferentially select forests that are ecologically distinct from unrepresentative regional averages. By digitizing and analyzing comprehensive offset project records alongside detailed forest inventory data, we provide direct evidence that comparing projects against coarse regional carbon averages has led to systematic over-crediting of 30.0 million tCO_2_e (90% CI: 20.5 to 38.6 million tCO_2_e) or 29.4% of the credits we analyzed (90% CI: 20.1 to 37.8%). These excess credits are worth an estimated $410 million (90% CI: $280 to $528 million) at recent market prices. Rather than improve forest management to store additional carbon, California’s offsets program creates incentives to generate offset credits that do not reflect real climate benefits.

**Significance Statement:** Forest carbon offsets are increasingly prominent in corporate and government “net zero” emission strategies, but face growing criticism about their efficacy. California’s forest offsets program is frequently promoted as a high-quality approach that improves on the failures of earlier efforts. Our analysis demonstrates, however, that substantial ecological and statistical shortcomings in the design of California’s forest offset protocol generate offset credits that do not reflect real climate benefits. Looking globally, our results illustrate how protocol designs with easily exploitable rules can undermine policy objectives and highlight the need for stronger governance in carbon offset markets.

## Introduction

Carbon offset programs issue credits to projects that purport to avoid greenhouse gas emissions or remove carbon dioxide from the atmosphere. When policymakers allow polluters to use offset credits to comply with policy requirements, these “compliance offsets” increase the quantity of greenhouse gas emissions allowed within a legally binding policy regime in exchange for climate benefits claimed somewhere else (1, 2). For example, an oil refinery that is subject to an emissions limit might purchase an offset credit issued to a forest owner who agrees to reduce or delay a timber harvest. The refinery can then claim the avoided forest emissions to compensate for its higher emissions. Compliance offsets have been widely used in cap-and-trade programs in the European Union and California (3, 4), to satisfy climate mitigation pledges made under the Kyoto Protocol (5), and, potentially, in the future implementation of the Paris Agreement (6).

Offsets are also controversial. Because compliance offsets enable higher emissions within legally binding policy regimes, they must reflect “additional” climate benefits that go beyond what is expected under counterfactual business-as-usual conditions (7). Compliance offsets’ additionality is therefore a fundamental prerequisite to their successful inclusion in climate policy, but this standard is not always achieved in practice. Prominent studies concluded that the world’s first carbon offsets programs, known as the Clean Development Mechanism (CDM) and Joint Implementation (JI), led to significant over-crediting from projects that made suspect claims about the additionality of their efforts or the plausibility of their emissions under counterfactual baseline scenarios (5, 8–11).

Because project-specific claims are hard to evaluate and easily exaggerated, some carbon offset programs, including the CDM, shifted to a second-generation or “standardized” approach. Under a standardized offset paradigm, offset protocols set common rules for determining project eligibility, setting projects’ baseline scenarios, and calculating the number of credits that should be awarded to eligible activities. Although standardized offset protocol rules help avoid suspect project-level claims, they also shift the risk of over-crediting from project-level claims to protocol-level calculations (4). One critical concern is the problem of adverse selection: prospective offset project developers know more than regulators about likely project-level baseline scenarios and have an incentive to preferentially select projects that naturally outperform regulators’ assumptions, potentially generating non-additional credits (12).

Thus, while a standardized protocol rule might prevent projects from customizing suspect methodologies to claim non-additional credits, that same rule might also introduce bias and create perverse incentives for project developers. Using a synthetic control analysis, for example, a recent study concluded that standardized baseline rules led to systemic over-crediting in REDD+ forest carbon offset projects in Brazil (13). Nevertheless, empirical evidence analyzing non-additionality and other kinds of over-crediting remains relatively rare because counterfactual scenarios are unobservable directly and can only be estimated indirectly through rigorous study with sufficient data and careful experimental design (10, 14).

Here, we analyze crediting errors from standardized baselines in California’s compliance offsets program, which plays a central role in the state’s prominent cap-and-trade program (4, 15–17). As of our study cutoff date of September 2020, the California Air Resources Board (CARB), which regulates the offsets program, had issued about 193 million offset credits across four different compliance offset protocols (18). These credits (each worth 1 tCO_2_e) represent about $2.6 billion as of recent market prices of $13.67/tCO_2_e (19).

## Results

Although California’s offsets program is open to many different kinds of projects, most credits come from a specific kind of forest offset project. About 82% of total credits are from CARB’s US Forest Projects protocol, which is open to application from forests anywhere in the continental United States and southern Alaska (20–22). Most forest offset credits come from “improved forest management” (IFM) projects, which claim to increase forest carbon storage through changes in forest management practices, such as increasing the length of timber harvest rotations. Critically, the bulk of credits issued to IFM projects are awarded “upfront” in projects’ initial reporting periods, based on the difference between initial on-site carbon stocks (as measured by field surveys) and the 100-year average carbon stock in projects’ baseline scenarios (as modeled by project developers according to standardized protocol rules) (16). Credits are also awarded annually for any increases in on-site carbon stocks due to forest growth, but the bulk of the credits in circulation — equal to about two-thirds of forest carbon offsets, and more than half of the entire carbon offsets program — come from upfront IFM credits (Figure 1A).

**Figure 1:**
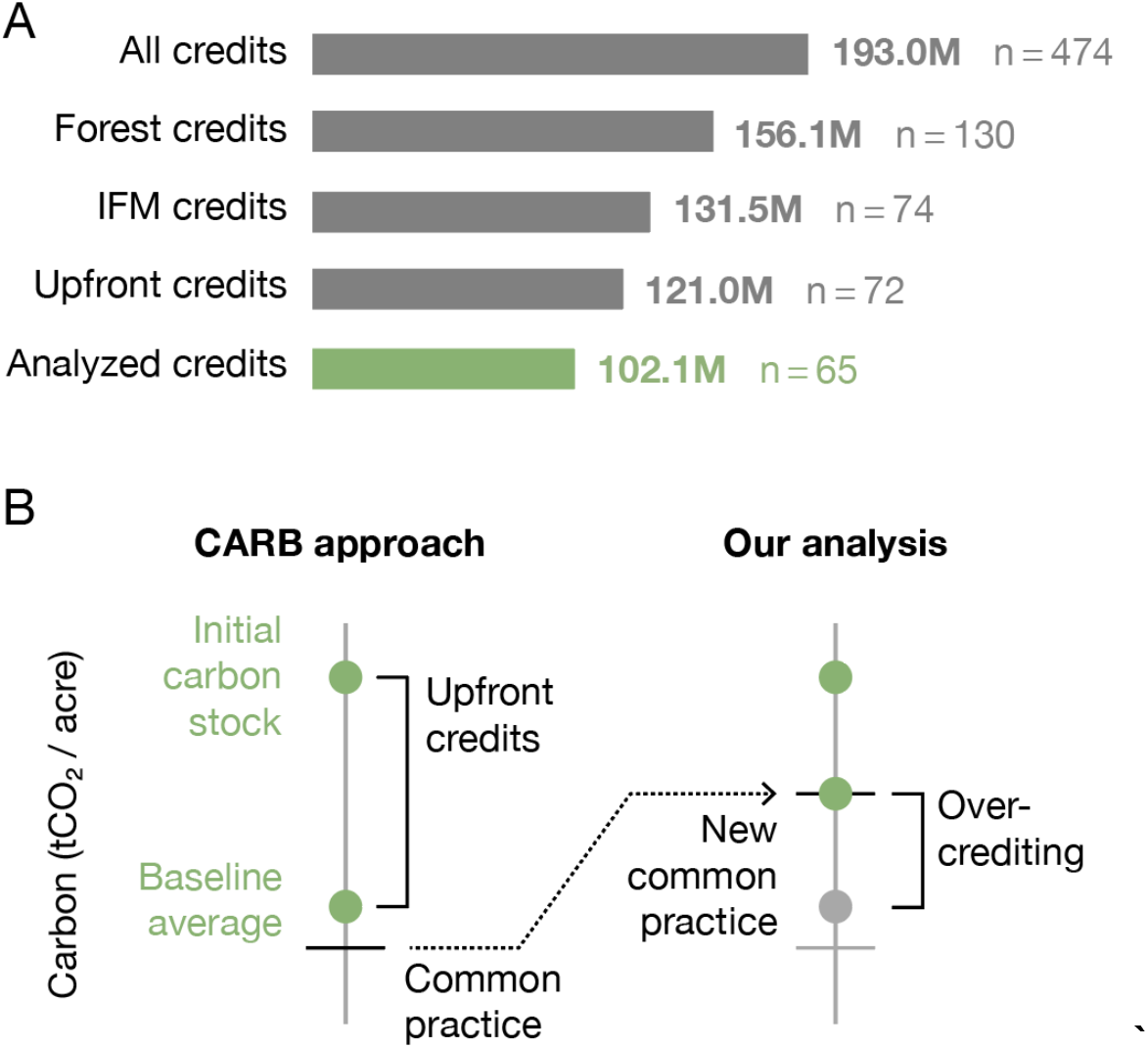
California’s offsets program. (A) As of September 2020, the California Air Resources Board (CARB) had issued 193 million offset credits, each worth 1 tCO_2_e, to 474 projects. The forest offsets protocol accounts for the vast majority of credits in the program, with most credits awarded to improved forest management (IFM) projects and most IFM credits earned in the form of initial, upfront credits calculated under standardized protocol rules. Limited public data disclosures restrict our analysis to 65 projects that earned 102.1 million upfront IFM credits, equivalent to about two-thirds of the forest offset program or about half of California’s total offsets program. (B) IFM projects are awarded upfront credits based on the difference between projects’ measured initial carbon stocks and the 100-year average carbon stocks projected in their baseline harvest scenarios. Under protocol rules, baseline averages must be equal to or greater than protocol-defined common practice calculations. Thus, erroneously low estimates of common practice can lead to over-crediting.

IFM project developers in California’s market have broad latitude to develop baseline scenarios, but cannot choose any baseline they like. Projects with higher-than-typical carbon stocks (72 of 74 of all compliance-period IFM projects) must report a baseline scenario with a 100-year average aboveground carbon stock that is no lower than “common practice” (Figure 1B). This rule prohibits projects from claiming they would harvest their forests below levels the protocol deems reasonable, defined as average regional carbon stocks from putatively similar forest types. IFM project baseline scenarios almost universally converge to common practice, a pattern that maximizes the number of upfront credits earned (Figure 2).

**Figure 2:**
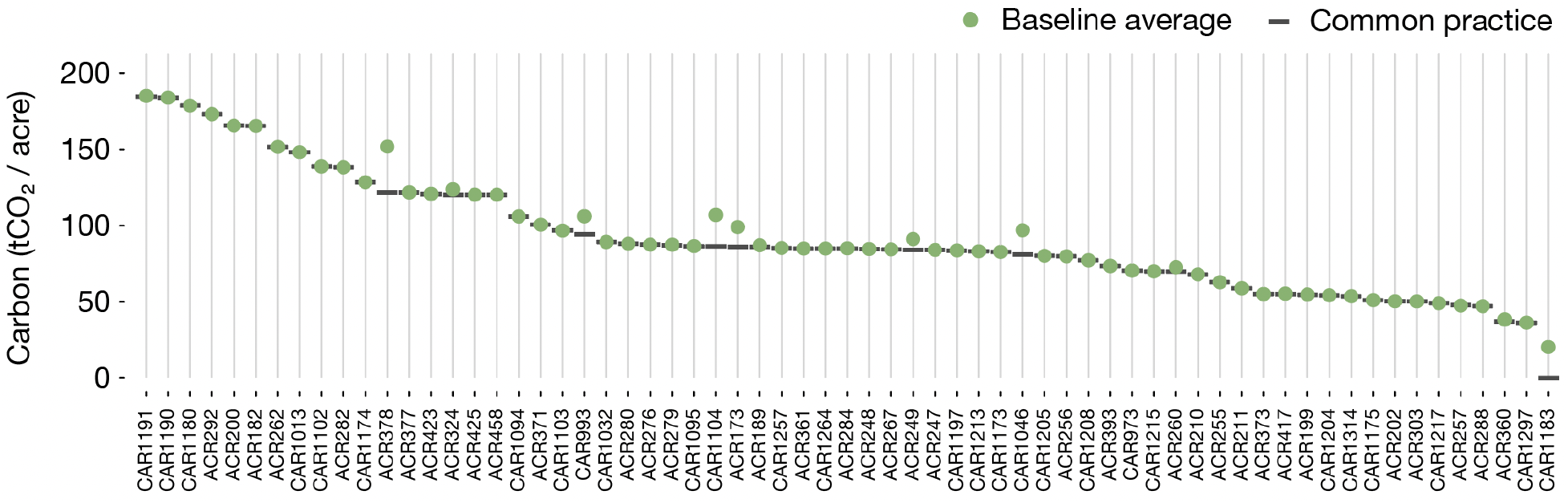
Forest carbon baseline scenarios converge to regional common practice estimates. Improved forest management (IFM) projects have baseline scenarios with 100-year average carbon stocks that converge on protocol-level calculations of regional common practice. The number of offset credits awarded to IFM projects depends on the difference between initial standing carbon stocks and the 100-year average carbon stock in IFM projects’ baseline scenarios, but these 100-year averages are constrained by protocol rules to be no lower than regional estimates of “common practice” for similar forest types. For each project, the green circle shows carbon in projects’ baseline scenario and the dark grey line shows common practice. 89% of projects analyzed are within 5% of common practice (mean Δ: 2.0 tCO_2_/acre, median Δ: 0.0 tCO_2_/acre).

As a result of these two features — a protocol rule prohibiting IFM projects’ average baselines from falling below common practice, and data indicating nearly all IFM projects report average baselines that converge toward or perfectly match common practice — the common practice numbers themselves are the primary determinant of upfront credits issued to IFM projects. Because upfront credits to IFM projects constitute the dominant share of all forest offset credits generated thus far (121.0 million credits, or about 77%) and a majority of all the credits in California’s entire offsets program (about 63%), the California regulator’s choice of common practice is arguably the single most important factor determining project crediting.

Common practice is calculated from the US Forest Service (USFS) Forest Inventory and Analysis (FIA) database, based on species combinations called “assessment areas”^1^ that span geographic regions termed “supersections” (23). These two concepts — assessment areas and supersections — were initially developed by the Climate Action Reserve (CAR), a nonprofit organization and carbon offsets registry (24). To construct supersections, CAR began with a set of eco-topographic regions called ecosections that were developed by the USFS to define management areas with similar geology, climate, and vegetation communities (25). CAR then combined ecosections together to create a novel set of supersections. Within each supersection, CAR defined one or more assessment areas to represent different species mixtures that are typical of forest types in that supersection. For example, CAR grouped various oak species within the Northern California Coast supersection into a single “Mixed Oak Woodland” assessment area, rather than considering each oak species individually. Finally, CAR used FIA data to establish common practice for each assessment area by taking the average carbon stocks of constituent forests. Thus, every supersection has one or more assessment areas, and each assessment area has a common practice estimate of average carbon stocks derived from FIA data across that assessment area’s supersection.

Although CAR initially developed these methods for the voluntary offsets market, the California regulator, CARB, subsequently adopted CAR’s methods for compliance purposes in its cap-and-trade program. CARB retained the same common practice numbers initially developed by CAR in CARB’s original 2011 US Forest Project protocol (20) as well as in a 2014 update (21). In a 2015 update, CARB worked with the USFS to update common practice numbers for the continental US and expand protocol eligibility to southern Alaska (22).

We developed a novel dataset from digitized public offset project records that enables direct estimates of crediting errors in California’s forest offsets program by comparing actual credits awarded against what would have been awarded using a more ecologically robust, project-specific determination of common practice. Instead of using a coarse regional average that combines ecologically distinct forest types into a single common practice, we estimate common practice from FIA plots that correspond to projects’ reported species composition (see Methods). We then re-calculate the number of credits projects would have received with our alternative and more appropriate estimates of common practice.

For many projects, our more ecologically robust estimate of common practice is higher than the supersection-wide values used in the California forest offsets program, which implies over-crediting. For a smaller number of projects we find a lower common practice, which implies under-crediting. To illustrate our results and make their causal factors concrete, we first describe in detail results for three representative projects (identified by their registry numbers ACR189, ACR361, and CAR1183) and then report aggregate statistics that show net over-crediting across the program as a whole.

Perhaps the most important example of over-crediting occurs in the Southern Cascades supersection, which ranges from the Pacific coast to the foothills of the Sierra Nevada and hosts the most offset projects of any supersection in California’s program. Within this region, CARB protocol rules specify that temperate, carbon-dense forest types like Douglas Fir (*Pseudotsuga menziesii*; average 122.5 tCO_2_e / acre) and Tanoak (*Notholithocarpus densiflorus*; average 192.4 tCO_2_e / acre) are averaged together with less-carbon-dense forest types that occupy more arid niches, like Ponderosa pine (*Pinus ponderosa*; average 60.4 tCO_2_e / acre). Comparing project carbon against this amalgamation of wet and arid forests causes projects like ACR189, which is composed primarily of Douglas fir (26% of basal area) and Tanoak (49% of basal area), to receive substantial upfront credits under protocol rules simply due to a mismatch between the species in the project and the species included in the regional average. By instead comparing ACR189 against FIA plots that contain primarily Douglas fir and Tanoak (see Methods), a more ecologically robust comparison, we estimate that ACR189 is over-credited by 135,869 tCO_2_e (90% CI: 85,481 to 185,917 tCO_2_e) or 50.1% of its total credits (90% CI: 31.5 to 68.6%).

Similar dynamics play out in the temperate rainforests of coastal Alaska, where orographically-induced precipitation and relatively warmer oceanside temperatures allow iconic species like Sitka spruce (*Picea sitchensis*; average 121.1 tCO_2_e / acre) and Western hemlock (*Tsuga heterophylla*; average 143.0 tCO_2_e / acre) to accumulate massive stores of carbon (26). ACR361, for example, consists of 94.9% Sitka spruce by basal area. Yet the common practice against which this Sitka-dominated forest is compared contains carbon estimates from far-less-carbon-dense forest types like Cottonwood (*Populus spp.*; average 41.4 tCO_2_e / acre) and Paper birch (*Betula papyrifera*; average 38.3 tCO_2_e / acre). Comparing ACR361 instead against other Sitka spruce forests from FIA measurements across the full coastal Alaska region indicates median over-crediting of 318,269 tCO_2_e (90% CI: −198,607 to 871,385 tCO_2_e) or 13.4% of its total credits (90% CI: −8.4% to 36.7%).

The most surprising example concerns a mixed conifer project, CAR1183, in the “sky island” forests of New Mexico (27). Despite the project consisting primarily of Douglas fir (37.1% of basal area) and Ponderosa pine (22.9% of basal area), the rules of the offset protocol allowed CAR1183 to enroll itself under the Pinyon (*Pinus spp.*) /Juniper (*Juniperus spp.*) Woodland assessment area. Perplexingly, in the 2011 and 2014 versions of the protocol, this assessment area had a common practice of 0 tCO_2_/acre (20, 21). Though CARB would later update this number to 8.74 tCO_2_/acre in its 2015 protocol (22), CAR1183 was developed under the earlier rules and earned 4.4 million upfront credits. In fact, under the earlier rules, any forest in that region would have been eligible for upfront credits. When more appropriately compared to FIA plots that contain Douglas fir and Ponderosa pine, CAR1183’s initial carbon stocks fall below the regional average. As a result, we estimate that 100% of the project’s claimed emission reductions are over-credited, a result that is robust across the full 5 to 95% confidence interval.

Across the program as a whole, we find evidence of systematic over-crediting (Figure 3). Of the 102.1 million tCO_2_e worth of upfront credits for which we have sufficient data to analyze, we estimate net over-crediting of 30.0 million tCO_2_e total (90% CI: 20.5 to 38.6 million tCO2e) or 29.4% of the credits we analyzed (90% CI: 20.1 to 37.8%). At recent market prices of $13.67 per offset credit (19), these excess credits are worth $410 million (90% CI: $280 to $528 million) – and likely more, as market prices would rise if market regulators took steps to correct for over-crediting.

**Figure 3:**
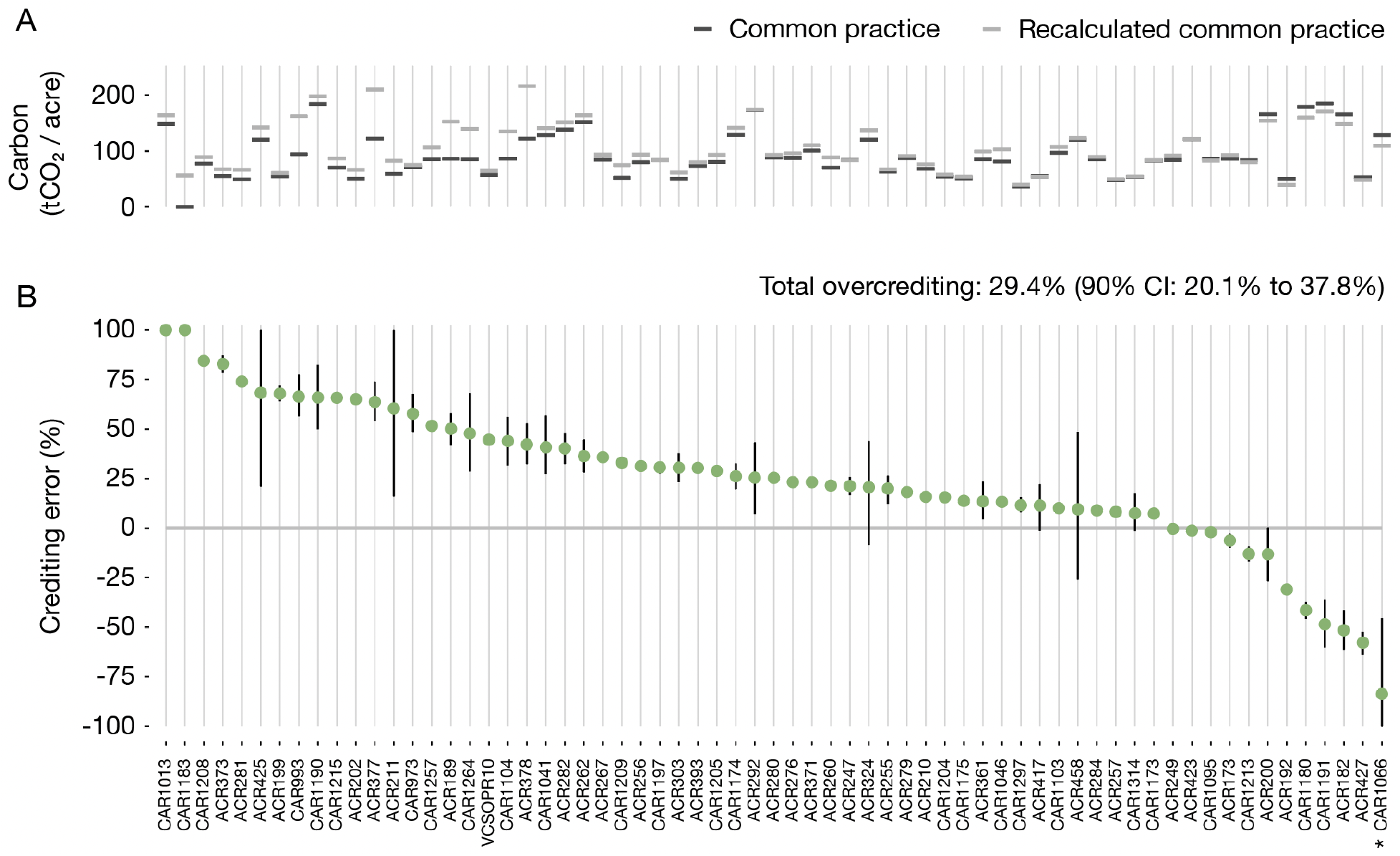
Estimated crediting error by project. We re-calculate the number of credits that would have been awarded to forest offset projects with a more ecologically robust measure of common practice. (A) The difference between common practice numbers used under protocol rules (dark grey lines) and our more ecologically robust common practice numbers (light grey lines) for each project. Over-crediting occurs when our common practice calculation estimate produces more carbon per acre compared to CARB’s common practice values, and under-crediting occurs when our common practice estimate results in less carbon per acre. (B) The extent of over- and under-crediting as a percentage of actual credits awarded to each project. Green circles indicate each project’s median estimate for over- or under-crediting, with vertical black lines spanning the 25th and 75th percentile estimates. Across the population of projects analyzed, total over-crediting is estimated at 30.0 million tCO_2_e total (90% CI: 20.5 to 38.6 million tCO_2_e) or 29.4% of the credits we analyzed (90% CI: 20.1 to 37.8%). (* Note that the bottom of the confidence interval for CAR1066 is truncated.)

Uncertainty ranges in our project-specific and program-wide results reflect uncertainty in the underlying USFS FIA data. Although CAR and CARB use point estimates of common practice, all calculations based on FIA data are subject to uncertainty. As indicated in Figure 3, some project-level estimates of crediting error have large confidence intervals (e.g. ACR211, ACR458) whereas others have narrow intervals (e.g. CAR1215, ACR260). The differences typically reflect the number of matching FIA plots in the project’s supersection (see Methods). Some locations have relatively few plots, which leads to higher uncertainties in estimates of common practice – notably in Alaska, where FIA sampling is sparse.

## Discussion

### Statistical bias in geographic regions

The fundamental challenge with awarding upfront offset credits via standardized protocol rules lies in defining an ecologically robust point of comparison. The California offsets protocol aggregates FIA data across assessment areas (species types) and supersections (geographic regions). We identify statistical patterns of project development that indicate widespread adverse selection, with projects preferentially located in forests where carbon stocks naturally exceed coarse, regional averages.

Part of the problem involves the way CAR and CARB construct supersections. The mixed conifer assessment area in the Southern Cascades supersection, which hosts more projects than any other supersection, provides a powerful illustration (Figure 4). The supersection is composed of three smaller USFS ecosections. Starting on the supersection’s western edge, ecosection M261B features relatively wet, carbon-dense forests with an average carbon stock for mixed conifers forest types of 150.5 tCO_2_/acre. But this ecosection is combined with two others, M261A and M261D, that have drier and less-carbon-dense forests (120.6 and 100.6 tCO_2_/acre, respectively). Under CARB’s protocol rules, the supersection-wide common practice for mixed conifer forests is 121.8 tCO_2_/acre, which makes an “average” forest in M261B immediately eligible for upfront credits. Although CAR and CARB both claim that combining ecosections with substantially different average carbon stocks does not change regional common practice by more than 10% (20, 21, 24), the creation of the Southern Cascades supersection appears to have violated this condition: the protocol’s 121.8 tCO_2_/acre is a −19% change from the M261B average of 150.5 tCO_2_/acre. Figure 4 shows clear clustering of projects within M261B, all of which likely take advantage of the ecologically suspect combination of ecosections.

**Figure 4:**
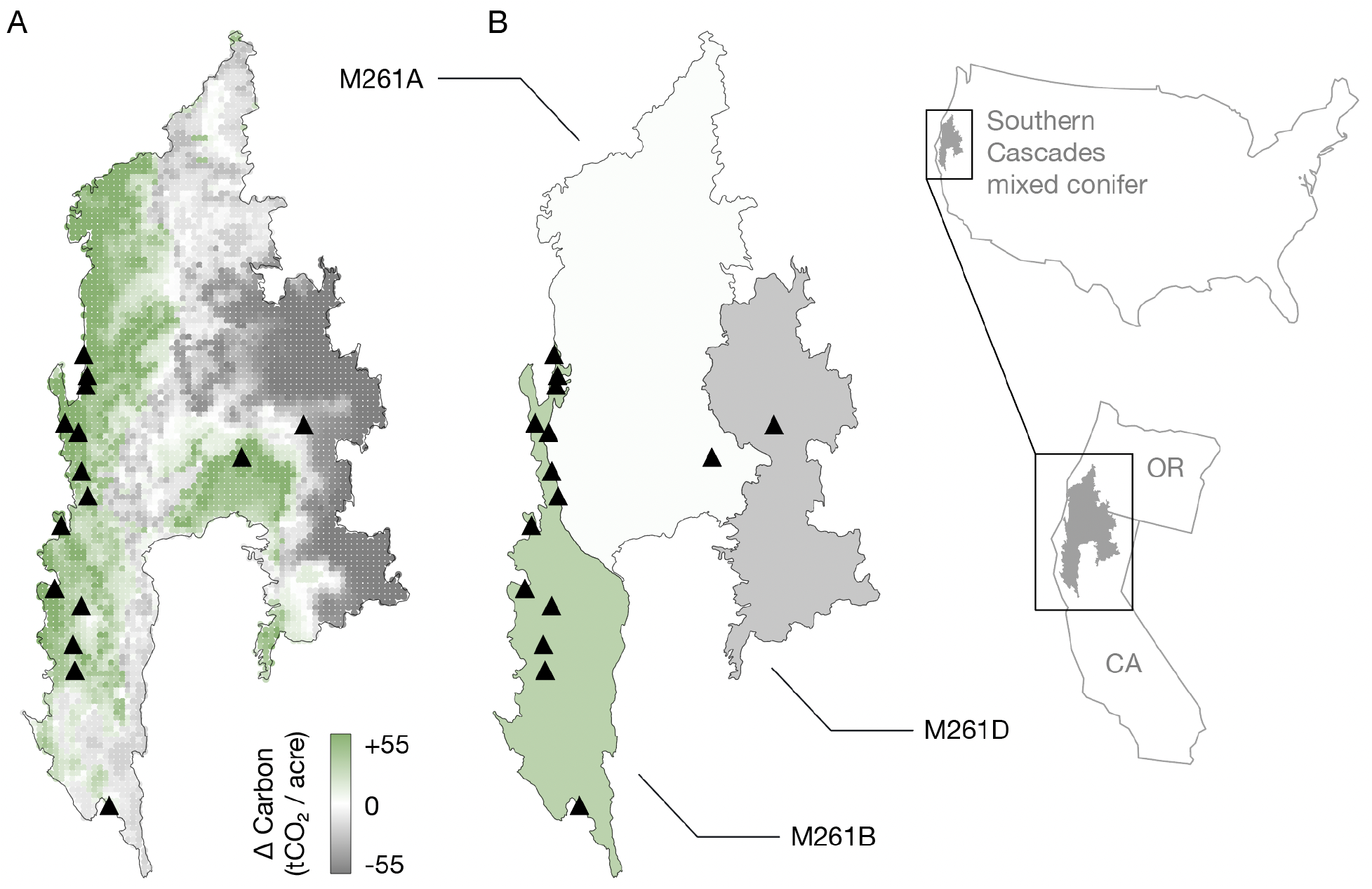
Arbitrage patterns in the Southern Cascades mixed conifer assessment area. One of the most extreme examples of over-crediting occurs in the mixed conifer assessment area of the Southern Cascades supersection. (A) The difference between standing live aboveground forest carbon in FIA plots that are climatologically similar to local conditions, and the supersection-wide average of all plots (see Supplementary Methods). Projects, represented with black triangles, cluster in carbon-rich areas, notably in wetter climates near the coast where carbon-dense forests grow. (B) The difference between ecosection- and supersection-wide common practice for mixed conifers. Three ecosections with distinct local carbon patterns were combined together to generate a supersection-wide common practice number that distorts ecological reality. The most carbon-rich ecosection (M261B) contains most of this supersection’s offset projects, which earn credits based on comparisons against supersection-wide averages that include dryer and less temperate ecosections (M261A, M261D).

The Southern Cascades supersection is an extreme example, but using any form of geographic aggregation introduces risks of adverse selection (28). Simple averaging over underlying variations in climate and its relationship to carbon storage necessarily introduces opportunities for adverse selection (Figure 4A). Biogeographers have long understood the challenge of drawing firm boundaries around ecological regions or categories of species because while boundaries help communicate with outside audiences, border regions are complex areas where the characteristics of separate regions interact (29–31). When used, spatial aggregation should be adopted carefully on the basis of ecologically meaningful boundaries and stress-tested for the potential to encourage adverse selection.

### Data limitations

Moving to species-specific analysis, such as our alternative approach to calculating common practice, partially addresses but does not completely avoid statistical challenges to a precise definition of common practice. Areas of the United States with extensive FIA sampling support common practice comparisons that are better grounded in ecology. But in other regions, notably Alaska, limited sampling is a barrier to robust estimates of common practice. The Alaska assessment area “North Coast Mountains, Chugach-St. Elias Mountains and Gulf of Alaska” has a mere 79 FIA plots, which serve as the basis for issuing over 9.5 million upfront credits. By contrast, the “Southern Cascades mixed conifer” assessment area in California and Oregon has upwards of 500 FIA plots. While the precision and uncertainty in our alternative estimate of common practice varies according to the rarity of forest types and prevalence of FIA data, the fact that our analysis accounts for variance in estimated carbon stocks across both species and space makes it more accurate and ecologically robust than the approach used in California’s program. Invoking the use of FIA data to assure the quality of a forest offsets program is not enough; a reliable protocol must also show how sampling density and statistical uncertainty are managed through rigorous protocol design (7).

### Baseline patterns and non-additionality

A key feature of our study is that it does not depend on counterfactual analysis to critique additionality claims. Claims that entire projects are non-additional are important to consider but difficult to evaluate quantitatively because counterfactual scenarios cannot be observed directly. In contrast, our analysis uses revealed program outcomes to directly estimate crediting errors. Nevertheless, the observation that nearly all offset projects choose baseline scenarios that converge on common practice (Figure 2) raises broader additionality concerns. It is possible that some projects’ “true” baseline scenario would be lower than protocol rules allow, such that converging on common practice would be appropriate for these projects. However, it is implausible that nearly all projects are in this situation, particularly since our re-estimate of common practice tends to be higher, not lower, than what the California program assumes. We also found evidence that projects specifically target common practice in baseline modeling. As one example, ACR373’s project documentation explains how linear optimization was used to drive the project’s baseline scenario as close to common practice as possible. Finally, we note that baseline over-crediting can be carefully combined with other estimates of over-crediting, such as extrinsic evidence that an entire project is non-additional or estimates of market-wide emission leakage effects, but we do not attempt that here.

### Policy implications

California law requires that offsets be “real, permanent, quantifiable, verifiable, and enforceable” (California Health & Safety Code § 38562(d)(1)) and that project baselines reflect “a conservative estimate of business-as-usual” conditions (California Code of Regulations, title 17, § 95972(a)(3)) (4). We estimate baseline over-crediting of 30.0 million tCO_2_e total (90% CI: 20.5 to 38.6 million tCO_2_e). One additional step is needed to evaluate the climate-equivalence claim made by California’s offsets program. The California forest protocol features a buffer pool, into which forest projects contribute a modest share of their total credits (up to about 20%) (32). The purpose of the buffer pool is to protect against risks to forest carbon from factors like fire, drought, and bankruptcy in order to ensure that forest carbon is stored for a 100-year permanence period, but credits in the buffer pool can, in theory, be used to compensate for any environmental inadequacy in the program. Our results indicate that over-crediting is likely larger than the program’s buffer pool, which contained 24.6 million tCO_2_e as of October 2020 (32). Even if over-crediting occurs at only the 5th percentile of our estimate (20.5 million tCO_2_e), addressing the environmental integrity of that outcome would deplete 83% of the buffer pool, leaving it severely undercapitalized in the face of growing climate risks (33, 34). This result calls into question whether California’s offsets program achieves the state’s policy goals.

## Materials and Methods

### Offset crediting components

Upfront credits in improved forest management (IFM) offset projects are awarded on the basis of differences between a project’s initial standing carbon and the 100-year average of aboveground carbon in its baseline scenario. Common practice constrains the minimum carbon in that baseline (Figure 2) and is computed separately for each supersection and assessment area. Supersections are geographic regions comprised of multiple ECOMAP 2007 ecosections (25). Assessment areas are groups of FIA forest types, each spanning a whole supersection, that are intended to reflect forest communities with similar ecological and economic attributes. Estimates of carbon from FIA are aggregated within each assessment area to derive common practice for that assessment area (20–22). Our analysis evaluates whether these aggregations lead to offset crediting errors.

### Digitized project records

We sourced project data from publicly available offset project data reports (OPDRs) submitted to CARB (see Supplementary Methods). We manually transcribed critical project attributes including total project acreage, initial carbon stocks, and the supersections and assessment areas involved in each project. We recorded 100-year average standing live aboveground carbon stocks in project baseline scenarios. For the initial reporting period, we recorded onsite carbon stocks (denoted IFM-1 and IFM-3) and the carbon stocks contained within wood products (IFM-7 and IFM-8), both for the baseline and project scenarios, as well as the project’s reported secondary effects and confidence deduction factors. We also transcribed all reported species with greater than 5% fractional basal area, on a per-assessment-area basis where data were available or else for the entire project. The schematized collection of records are available at http://dx.doi.org/10.5281/zenodo.4630684.

### Verification of crediting calculations

We verified the accuracy of our digitization by replicating actual project crediting calculations directly from project data, using Equation 5.1 from the 2015 CARB US Forest Projects protocol (22). Two members of our project team independently performed this exercise to ensure quality and converged on a unified result. We compared these estimates to the CARB-reported project issuance table dated September 9th, 2020 (R^2^=0.998; see Figure S1) (18).

### Forest inventory data

We analyzed data from the Forest Inventory and Analysis (FIA) database using rFIA, an open source software package that implements statistical practices recommended by the US Forest Service (35, 36). We developed queries to estimate the total aboveground carbon and total acreage for every supersection, assessment area, site class, inventory period, and forest type, along with their variances. All of our subsequent estimates of common practice (either using CARB’s approach or our alternative) sum carbon and acreage separately, before taking the ratio to report tCO_2_/acre (35, 37).

### Verification of common practice

CARB’s reported common practice aggregates carbon across all forest types within each assessment area on a supersection-wide basis. We confirmed that, from our processing of FIA data, we could independently reproduce CARB’s common practice values by comparing our estimates directly to the values reported in the CARB-provided Assessment Area Data File described in the forest offset protocol and available on CARB’s website (R^2^=0.97, RMSE=4.94 tCO_2_/acre; see Figure S2A) (22).

### Alternative species-specific common practice

We developed an alternative, more ecologically robust definition of common practice using project-reported species composition data. We compare each project against a project-specific (and therefore more representative) subset of FIA data, as opposed to the default, coarse regional averages of the CARB protocol. We built a classification algorithm (as described below) to map species composition (as reported in project OPDRs) to forest types (a set of canonical species groupings reported by FIA). For every project, the classifier returns a list of forest types and the probability that the project belongs to those forest types. We then use these forest-type assignments to estimate common practice from FIA plots that share those forest type codes.

### Classification algorithm

We fit a radius-neighbors classifier on a per-condition basis using pairs of two reported quantities in the FIA database: fractional basal area per species (derived from per-tree measurements) and recorded forest type code. Intuitively, the classifier takes species composition data as an input and estimates the probability of that species mixture belonging to different FIA-defined forest types based on relative similarity to the species composition of FIA plots. We fit a separate classifier for each supersection, based on all FIA plots within the supersection boundaries. We used grid search and 5-fold cross-validation to find the radius (“neighborhood”) that maximized the classifier’s ability to predict FIA-reported forest types from FIA-observed species data. The median, weighted F1 accuracy score (which considers Type I and Type II classification errors) across all classifiers was 0.78, with 1 being the best score (see Supplementary Methods).

### Calculation of over- and under-crediting

We use our alternative species-specific common practice to calculate a new 100-year average carbon stock in each project’s baseline scenario, assuming that the new common practice would constrain average baseline carbon stocks. Rather than replace the common practice reported by the project with our estimate, we scale a project’s reported common practice by the assessment-area-weighted ratio of our alternative calculation of common practice to our own re-calculation of CARB’s assessment area estimates (Figure S2A). Scaling by this ratio ensures that changes in common practice are due exclusively to changing assumptions about how FIA data is aggregated (see Supplementary Methods). These steps allow us to estimate the credits that would have been awarded to actual projects using our alternative common practice calculation. We obtained confidence bounds on our estimates of crediting error through Monte Carlo error propagation. Using variances of carbon per acre from FIA for each forest type and assuming gaussian noise, we sampled 1000 random draws of FIA carbon estimates and on each draw calculated the crediting error for individual projects. Throughout, we report the 5th, 50th, and 95th percentiles of the resulting distribution.

## Author contributions

G.B., D.C., J.F, and J.J.H. designed the research; G.B. digitized the project report data; G.B., D.C., J.F, J.J.H., and B.H. performed the research and analyzed the data; all authors contributed to interpreting the results and writing the paper.

## Acknowledgements & Funding

We thank Olaf Kuegler of the USFS for feedback on the statistical approaches needed to translate FIA data into the common practice values used by CARB. Hunter Stanke provided critical feedback on the use of rFIA. Jim Omernik helped in understanding the history of the USFS ECOMAP project, as well as his own work and the separate efforts of Bob Bailey to draw maps capturing the distinct biogeography of American forests. Thanks to Matthew D. Potts, David G. Victor, Cindy Chiao, Oriana Chegwidden, and Freya Chay for comments on a draft manuscript. Any errors are the authors’ sole responsibility.

G.B. acknowledges support from the Black Rock Forest Postdoctoral Fellowship in Forest Ecology. CarbonPlan acknowledges support from Microsoft AI for Earth for the digitization of offset project records. W.R.L.A. acknowledges funding from the David and Lucile Packard Foundation. A.T.T. acknowledges funding from NSF Grant 2003205.

The authors declare no conflicts, financial or otherwise, that could be perceived as influencing the research described here. D.C. is the Vice Chair of California’s Independent Emissions Market Advisory Committee, but does not speak for the Committee here.

## Supplementary Methods

Here we present key sections of our methods in more detail. Section names below correspond to matching sections of the Brief Methods in the primary article.

### Digitized project records

We sourced project data from project-submitted “offset project data reports” (OPDRs), the official documentation offset projects submit to CARB. These documents are made available by the three offset project registries that help CARB administer California’s offset program: the <u>Climate Action Reserve</u> (CAR), the <u>American Carbon Registry</u> (ACR), and <u>Verra</u> (VCS).

For each project, we transcribed project details described in the “initial” and “annual” OPDRs. In the rare case where initial and/or annual OPDRs were unavailable, we sourced information from the project’s listing information (which is also hosted by the offset registries), taking note of the discrepancy. We recorded critical project attributes such as total project acreage, reported initial carbon stocks, and the supersections and assessment areas of each project. For project baseline scenarios, we recorded the 100-year average standing live aboveground carbon stock. In most cases, this variable was directly reported in the text of the initial OPDR and its supplements. However, in some cases, only a graphical depiction of the baseline scenario was provided. In these cases, we used a <u>graph digitization tool</u> to infer 100-year average standing live aboveground carbon. In some cases, when both common practice and the 100-year average standing live aboveground carbon stocks were both clearly displayed and were visually indistinguishable, we recorded the 100-year average standing live aboveground carbon stock as being equal to common practice.

For the initial reporting period, we recorded onsite carbon stocks (denoted IFM-1 and IFM-3) and the carbon stocks contained within wood products (IFM-7 and IFM-8), both for the baseline and project scenarios as well as for the project’s reported secondary effects. Onsite carbon stocks (IFM-1 and IFM-3) for the “project scenario” were further adjusted by the project-reported confidence deduction factor that reduces projects’ earned offset credits due to statistical uncertainty in on-site carbon measurements above a 5% threshold (1). These data allowed us to recalculate the number of ARBOCs that should have been granted to the project based on publicly available documents for each projects’ first reporting period.

When reported, we also transcribed details about the species composition of each project. As detailed in Section 3.1(a)(1) of the 2015 US Forest Projects protocol, projects must report the species makeup of each individual assessment area in terms of fractional basal area, which is then compared against an assessment area specific “Species Diversity Index” that is reported alongside common practice numbers in the CARB-provided Assessment Area Data File. We recorded all species, on a per assessment area basis, with greater than 5% fractional basal area. Some projects only reported species composition on a whole-project basis, which we recorded and, in subsequent analyses, assumed all assessment areas had that same, fixed species composition (see below). All species were denoted using the appropriate FIA species code from Appendix F of the FIA User Guide (2).

While not used directly in our analysis except for plotting purposes, shapefiles for projects were obtained from the California Air Resources Board’s online Credit Issuance Map, accessed at https://webmaps.arb.ca.gov/ARBOCIssuanceMap/ using the ArcGIS MapServer API. A archival copy of standardized and processed shapefiles as GeoJSON is available at http://doi.org/10.5281/zenodo.4630684.

Our full digitized database can be downloaded in JSON and CSV formats. Most of the associated metadata, alongside a subset of our analysis results, can be browsed in an interactive map, all available at https://carbonplan.org/research/forest-offsets. A sample project record from the database is shown in Supplementary Excerpt 1. Archived versions of all primary source materials (e.g. ODPRs) are available in a Zenodo archive (3).

**Supplementary Excerpt 1.**
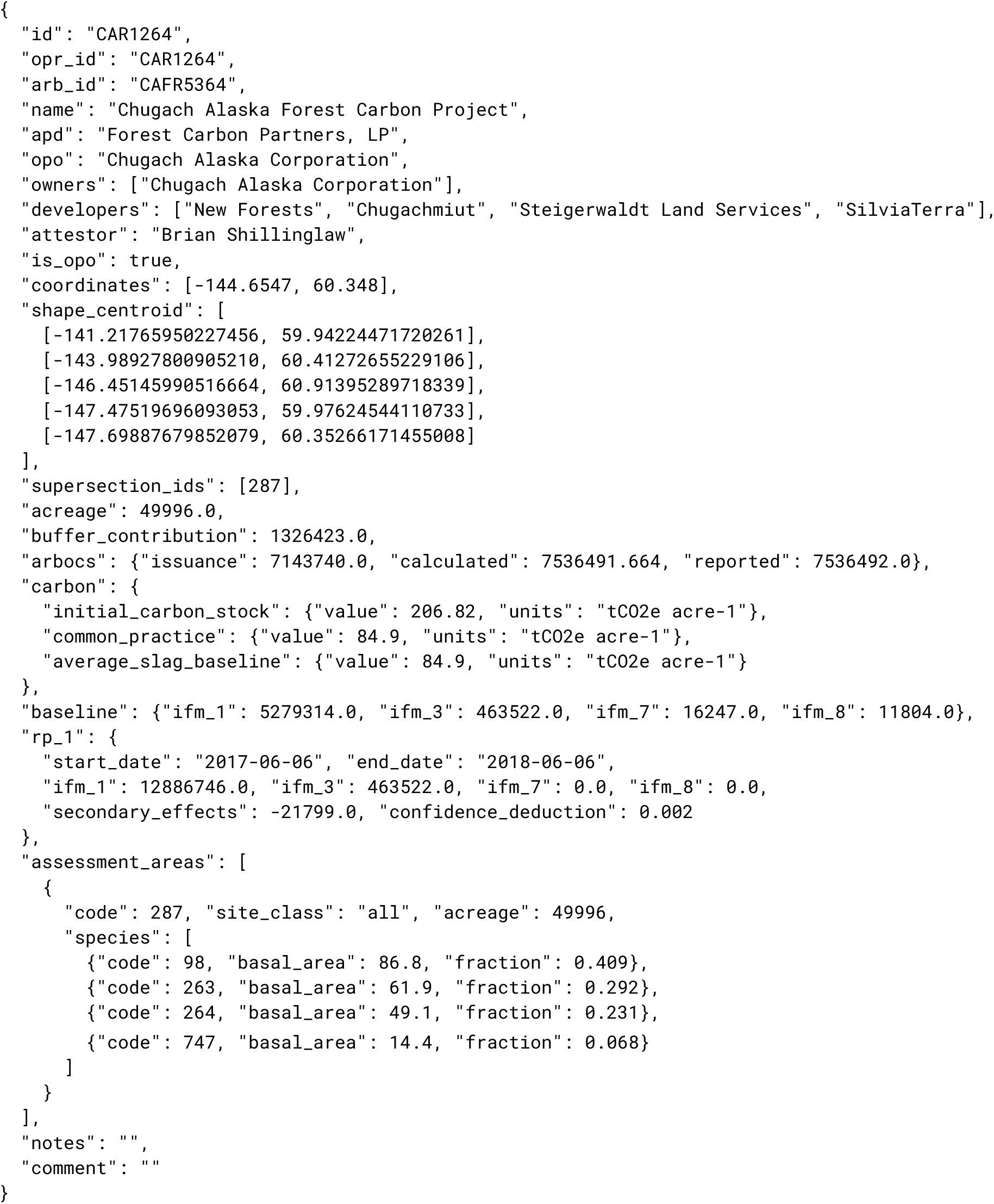
Sample project record (CAR1264) from the database.

Our final digitized database contains 93 entries, representing all credited IFM offset projects we were able to identify that were credited as of the 2020-09-09 CARB issuance table (4). We identified 19 of those projects as having participated in the CARB Early Action (EA) program phase and subsequently “graduated” into the compliance program. Reporting details about the first reporting period of these projects required examining far less standardized “project design documents” (PDDs, as opposed to OPDRs). In some cases, project details from the EA project, as reported in the PDD, differed from the values reported in the graduated project’s OPDR, raising further concerns of data consistency. Given the less standardized project documentation, combined with the fact that many Early Action projects were initiated under a slightly different set of rules than the final 2011 CARB US Forest Projects protocol, we opted to exclude all Early Action projects from our analysis so as to ensure we applied the same data entry and analysis methods to all projects. This decision ensures that any rule changes between the EA and the compliance program do not influence our results.

Our primary analysis focused on the 74 remaining projects that entered the CARB offset protocol under the finalized rules of one of the 2011, 2014, and 2015 US Forest Projects protocols. Of those 74 projects, 72 projects received “upfront” offset credits due to the project’s initial carbon stocks exceeding protocol-determined common practice. Of those 72 projects, 65 projects could be analyzed using the species classification approach described below. For five of the unanalyzed projects, we were unable to identify a list of species in any publicly available documents (projects ACR248, ACR288, CAR1094, CAR1217, and CAR1032). One project (CAR1102) reported species composition for the entire project, as opposed to per assessment area. Under typical circumstances, our method uses the project wide species composition to estimate standing carbon for each assessment area. However, in this case, CAR1102 spans two supersections and one supersection (Northern California Coast) did not have observations for some of the oak forest types present in the project. This missingness results in the inability to estimate standing live carbon for over 25% of the project’s basal area. Rather than make assumptions about how species map to various supersection/assessment area combinations, we excluded the project from consideration. Finally, ACR360, a project in Alaska’s Copper River Basin falls entirely within the USFS ecosection 133B. However, there are no FIA plots in ecosection 133B, so we did not include the project in our analysis.

### Verification of crediting calculations

We verified the accuracy of our digitization by replicating actual project crediting calculations directly from project data, using Equation 5.1 from the 2015 CARB US Forest Project protocol (1). We used the September 9, 2020 version of the California Air Resources Board’s Credit Issuance Table as the official record of how many ARBOCs were awarded to each IFM project (4). CARB updates its official issuance table on a bimonthly basis (https://ww2.arb.ca.gov/our-work/programs/compliance-offset-program/arb-offset-credit-issuance).

**Supplementary Figure 1.**
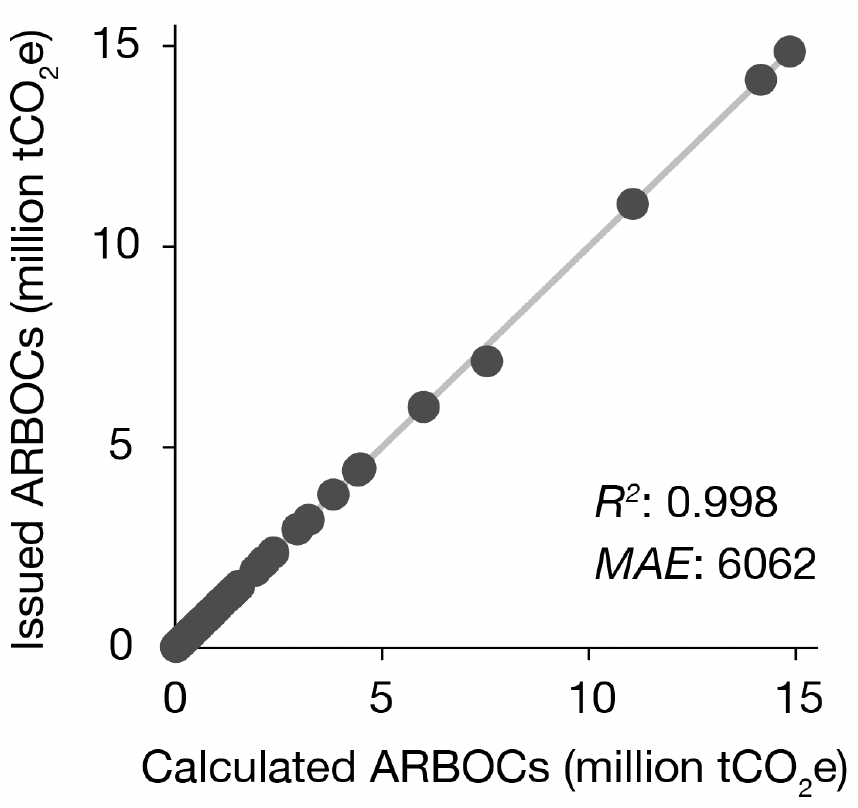
Comparison between ARBOC_Calculated_, our rederived calculation of ARBOCs awarded to each project from data contained within project OPDRs, and ARBOC_Issuance_, the actual number of ARBOCs awarded to a project by CARB. Mean absolute error of 6062 tCO_2_e.

Despite obtaining extremely similar results (Figure S1), we identified some small differences, which we describe comprehensively in an Appendix at the end of this document. For clarity, we introduce three pieces of notation to distinguish various offset credit (ARBOC) estimates: ARBOC_Issuance_ refers to ARBOCs issued by CARB and reported in the issuance table, ARBOC_Reported_ refers to ARBOCs reported by the final in their OPDRs, and ARBOC_Calculated_ refers to ARBOCs calculated in our analysis, based on the data reported in project OPDRs. Two members of our project team (G. Badgley and B. Haya) independently performed this exercise to ensure quality and converged on a unified result. In some instances, we refer to their findings by name. To our knowledge, this work reflects the first public attempt to audit project reported ARBOCs.

### Forest inventory data

We precomputed estimates of above ground live carbon from the USFS Forest Inventory and Analysis (FIA) database using the rFIA package, an open source software package that implements the queries necessary to replicate USFS statistical procedures (e.g., expansion factors, stratum weighting) for deriving robust inferences from FIA survey data (5).

For every assessment area within every supersection, we calculated above ground live carbon for each forest type code (FORTYPCD) using the ‘biomass’ function from the rFIA software package. Specifically, within rFIA, we loaded data from all US states overlapping the given supersection, matched inventories across states, and removed any samples falling outside the geographic boundary of the supersection. We then calculated above ground live carbon for all accessible, forested conditions (COND_STATUS_CD=1) on private land (OWNGRPCD=40). Finally, CARB common practice estimates for states in the Pacific Northwest work unit (AK, CA, OR, WA) used regional biomass estimates, as opposed to using biomass as reported in the default TREE table (O. Kuegler, personal communication). For these four states, we used regional biomass estimates reported in the TREE_REGIONAL_BIOMASS table. We included these values by setting DRYBIO_SAPLING, DRYBIO_WDLD_SPP, and DRYBIO_TOP equal to zero, retaining DRYBIO_STUMP, and replacing DRYBIO_BOLE with the reported per-tree value of REGIONAL_DRYBIOT from the TREE_REGIONAL_BIOMASS table.

When supersections spanned multiple states we harmonized FIA evaluations across all states using the rFIA function clipFIA, with option matchEval set to TRUE. We subset data spatially using the ‘polys’ argument in ‘biomass,’ meaning that plots were assigned to as supersection based on the “fuzzed,” publicly reported latitude and longitude values. We used the temporally indifferent method (“TI”), meaning our standing live carbon estimates pool together all FIA survey panels within a single inventory period. Whenever possible, we reported the carbon estimates as the median of inventories ending between 2010 and 2013, so as to be consistent with the snapshot of FIA data used by CARB to produce its own estimates of common practice. In the rare cases where no inventory period ended between 2010 and 2013, we took the median of all inventories from 2013 onward.

These queries yielded a point estimate and variance for above ground carbon and forested area for each forest type code and inventory. These estimates provide the inputs into our subsequent analyses.

### Verification of common practice

Given an estimate of carbon for each forest type, we can estimate different versions of common practice by aggregating in different ways. To validate our use of FIA data, we first used our carbon estimates to compute common practice as computed by CARB.

**Supplementary Figure 2.**
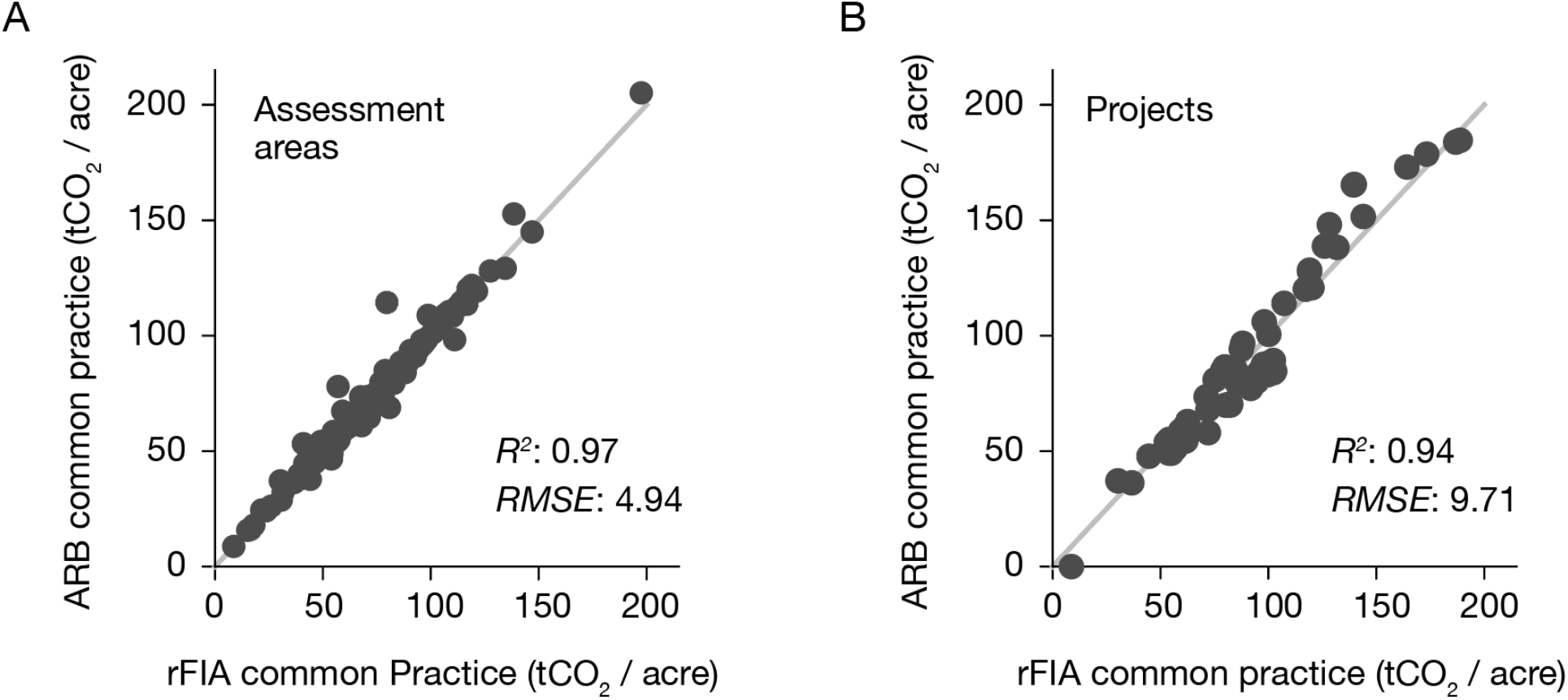
Comparison of common practice per assessment area (A) and on a per project (B) basis.

First, for each assessment area, we aggregated our carbon estimates within the assessment area, and compared the result directly to the value reported by CARB in 2015. Across all assessment areas containing projects, we found extremely high similarity (R^2^=0.97, RMSE=4.94 tCO_2_e/acre). We limited our comparison to supersections containing credited projects.

Second, for each project, we used the project-reported fractional decomposition by assessment area to compute a weighted average for the project (CP_0_), and compared these to common practice as reported by individual projects in project documentation (CP_ARB_). We found high agreement (R^2^=0.94, RMSE=9.71). On average, our estimates (CP_0_) were 3.2% higher than CARB’s reported values (CP_ARB_).

Minor deviations in both cases could be due to differences in exact inventories used as well as revisions to the underlying data and stratifications. Because FIA data archives can be updated, the newer version of the FIA database we use might have slightly revised data as compared to the database used by CAR and CARB. In particular, projects were registered under three different versions of CARB’s US Forest Project protocol that were issued in 2011, 2014, and 2015. The handling of a concept called site class in the protocol changed in 2015, and our estimates use the approach CARB employs in its 2015 protocol.

As outlined in the Brief methods and described in detail below, these small differences are highly unlikely to influence our analysis of over- or -under-crediting because we calculate *proportional changes* in common practice, each derived from the same underlying data, thereby isolating the effect of how FIA data is aggregated to calculate common practice, as opposed to uninformative differences between our estimates of common practice and the FIA values used by in the CARB US Forest Project protocol.

Together, these results validate our ability to compute common practice from FIA data, and thus allow us to consider variants of common practice calculated using alternate aggregations.

### Classification algorithms

Our analysis of over- and under-crediting relies on an alternate method of calculating common practice based on the species-specific composition of each project. This calculation relies on a radius-neighbors classifier that maps the species composition (as reported in project OPDRs) to forest types (as reported by FIA on a per-condition basis). Here, we describe that algorithm in detail.

Intuitively, the classifier takes as input the fraction of each species, and produces as output the probabilities of it belonging to one of several forest type codes. We implemented the classifier using the ‘RadiusNeighborsClassifier’ method from scikit-learn (6). Rather than look for n-nearest neighbors, the ‘RadiusNeighborsClassifier’ produces a classification estimate based on all training data that falls within a fixed radius of the observation. This approach is useful when classifying observations within potentially sparse “neighborhoods.” To train the classifier, we used pairs of two observed quantities on a per-condition basis — fractional basal area per species and recorded forest type code. We trained a separate classifier for each supersection. Grid search was used to find the radius that maximized performance with 5-fold cross-validation. Supplementary Table 1 reports the weighted F-1 accuracy scores of the final models, as evaluated on a 20% hold-out sample. F-1 scores are the harmonic mean of classifier recall and precision 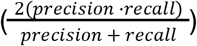 with a score of 0 being the worst score possible and 1 being the best.

After training the classifier, for each assessment area within each project, we used the reported species composition to estimate a forest type code distribution, and used that distribution in our alternate common practice calculation. Although most projects report species composition in terms of a per-species fractional basal area for each assessment area, 24 projects instead report species composition for the entire project. In these cases, we used the classifier as above, assigning the whole-project species composition to all assessment areas.

As a final check, we screened classifier performance by comparing the project’s species list against the classifier outputs using the outputs shown in Appendix 2.

**Supplementary Table 1.** Weighted F1-scores for the per supersection classifiers. F1-scores provide a weighted average of classification recall and precision, with 1 being the highest possible value.

| Supersection | Weighted F1-Score |
| --- | --- |
| Gulf Coastal Plain | 0.74 |
| Northern California Coast | 0.75 |
| Southern Cascades | 0.82 |
| Laurentian Mixed Forest Northern Highlands | 0.77 |
| Adirondacks & Green Mountains | 0.74 |
| St Lawrence & Mohawk Valley | 0.76 |
| Okanogan Highland | 0.88 |
| Columbia Basin | 0.89 |
| White Mountains - San Francisco Peaks - Mongollon | 0.93 |
| Lower New England - Northern Appalachia | 0.73 |
| Central New Mexico | 0.94 |
| Northwest Cascades | 0.87 |
| Allegheny & North Cumberland Mountains | 0.69 |
| Southern Allegheny Plateau | 0.68 |
| Laurentian Mixed Forest Southern Superior | 0.79 |
| Eastern Broadleaf Forest Cumberland Plateau | 0.69 |
| White Mountains | 0.78 |
| Maine - New Brunswick Foothills and Lowlands | 0.78 |
| Southeast and South Central Alaska | 0.91 |

### Calculation of under- and over-crediting

Our analysis of over- and under-crediting considers three versions of common practice: the common practice reported by each project (CP_ARB_), a calculation of common practice meant to be as comparable as possible to the approach used by CARB used (CP_0_) by aggregating within assessment areas, and a recalculation of common practice using the species classification method described above (CP_1_).

To calculate CP_1_, for each project we use the probabilities returned by the classifier to compute a weighted average of tCO_2_/acre across forest types. This approach aggregates over only the forest types that match the species composition of the project and are within the geographic bounds of the supersection, as opposed to uniformly aggregating over a discrete list of forest type codes in a predefined assessment area, which may not correspond to the actual species in the project. For example, our approach prevents projects that are primarily Douglas Fir to be classified as Pinyon/Juniper (e.g. CAR1183), and it prevents Douglas Fir projects from being compared to an aggregation of Douglas Fir and Ponderosa Pine (e.g. in the Southern Cascades Mixed Conifer assessment area).

Ideally, we would compare our classification result (CP_1_) to the actual common practice reported by projects (CP_ARB_). However, several factors make this comparison potentially problematic. First, projects have been developed under two different sets of common practice rules, depending on whether projects were developed before and after the adoption of the 2015 Forest Offset Protocol (FOP). It is especially difficult to recreate how the 2011 and 2014 FOP common practice values treated “site class” in their calculations. Prior to the 2015 revision, the cutoff between “high” and “low” site class varied from supersection to supersection (and perhaps even from assessment area to assessment area), making recreating the earlier common practice values exceedingly difficult.

While it is particularly difficult to identify how calculations before 2015 were performed, we have reason to think they relied at least in part on incomplete data, as evidenced by the fact that there were assessment areas with a common practice of 0, which is biologically impossible. Second, and related, we know our analysis does not use the same version of FIA that was used to compute all instances of CP_ARB_, because FIA data are updated and changed over time. Furthermore, we could find no public documentation of the data or code used to calculate common practice prior to the 2015 version of the CARB protocol.

Because of these possible sources of error, we avoid comparing CP_1_ directly to CP_ARB_. Instead, we focus on the sensitivity of common practice calculations to assumptions about aggregation, which ought to be comparable across projects and across time. We can directly calculate that sensitivity by comparing our estimate of CP_1_ to our estimate of CP_0_, both of which are derived from the same preprocessing of the same underlying FIA data. Calculating sensitivity this way isolates the effect on common practice of changing assumptions about aggregation. Given that our estimates of CP_0_ are highly similar to CP_ARB_, we can confidently use this sensitivity to infer potential under- and over-crediting.

Having calculated the ratio of CP_1_ to CP_0_, we calculate a new common practice for each project by using this ratio to rescale the project’s actual common practice.

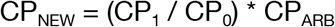

We then recalculate the CARB offset credits (ARBOCs) that would have been awarded to the project on the basis of CP_NEW_. Recall that, under the protocol, awarded upfront credits are based on the difference between the IFM-1 project scenario and IFM-1 baseline scenario. Further, the IFM-1 baseline scenario is constrained to be above common practice, and empirically, nearly all projects present a baseline that is at or only slightly above common practice (89% of projects within 5% of common practice). To calculate potential over- or under-crediting, we assume that new IFM-1 baselines would similarly be set to this new common practice. A caveat is that IFM-1 incorporates both above and below ground carbon, which is calculated in the protocol using per-species allometric equations, whereas CP_NEW_ only considers above ground carbon. To correct for this difference, for each project, we estimated the ratio of above ground to below ground carbon by dividing IFM-1 in the project scenario by the initial above ground carbon stock reported by the project, which typically yields a scale factor slightly greater than 1 (1.23 +/− 0.04 mean/sd).

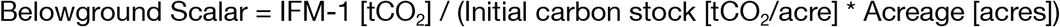

We multiply CP_NEW_ by project acreage and the scale factor, and set this value as IFM-1 in the baseline scenario. Given a new baseline IFM-1 for a project, we can then recalculate ARBOCs using Equation 5.1 of the 2015 US Forest Projects protocol.

We express under- and over-crediting in units of million tCO_2_e and also as a percent of the total number of offset credits issued to the project. We also sum under- or over-crediting across projects and express this sum as a fraction of the total ARBOCs of all projects analyzed. We use ARBOC_Calculated_, as opposed to ARBOC_Issued_, to account for the fact that details provided in the digitized records occasionally differ from the documents used by CARB for issuance.

Note that IFM-3 (standing dead carbon) is ignored in this analysis. For the majority of projects (54%), IFM-3 in the baseline scenario and project scenario are equal, suggesting that this is not a major source of credits. We are thus not estimating any over- or under-crediting for IFM-3.

Note also that any systematic bias in our estimates of CP_0_ relative to CP_ARB_ could potentially overestimate (or underestimate) our re-crediting calculations. Specifically, if we systematically overestimated CP_0_, then we underestimated over-crediting; similarly, if we systematically underestimated CP_0_, then we overestimated over-crediting. As reported above, our estimates of CP_0_ are well matched to CP_ARB_ (R^2^=0.94, RMSE=9.76), and on average were 3.2% higher than CP_ARB_. If anything, the fact that we overestimate CP_ARB_ likely makes our overall finding of net over-crediting conservative. In addition, we found no evidence for a systematic relationship between error in our estimate of CP_0_ and our estimates of crediting error (r=0.06).

We used Monte Carlo error propagation to bound our estimates of crediting error. Using variances of total carbon per acre as reported by rFIA, and assuming gaussian noise, we sampled 1,000 random draws of FIA carbon estimates for CP_1_ and on each draw calculated crediting for individual projects and across the full portfolio of projects. We use these distributions to report 5th, 25th, 50th, 75th, and 95th percentiles for our estimates of crediting error.

In general, variability in our estimates of crediting error was largest when the number of FIA conditions available for analysis was small.

### Special methods for CAR1183

In one unusual case, CAR1183, we had to slightly modify our primary method due to a factual error in CARB’s 2011 and 2014 forest offset protocols. Because we had to change our method for this project, we performed an additional and complementary analysis to evaluate the robustness of our results.

When CAR1183 was initially listed, the entire project was assigned to the Central New Mexico Pinyon/Juniper assessment area, which CARB assigned a common practice (CP_ARB_) of 0 tCO_2_e per acre — a clear error, as this number implies forests in the region contain no CO_2_. Weeks after the project was listed, CARB’s 2015 US Forest Projects protocol fixed this error by (i) updating the Central New Mexico Pinyon/Juniper assessment area to have a non-zero common practice and (ii) introducing a “Mixed Conifer” assessment area to the supersection.

Despite these revisions, the fact that the project’s reported CP_ARB_ (see Extended Methods Equation 1) equaled zero means that our estimate of CP_NEW_ would always equal zero. This because our method multiplies the ratio of rFIA derived common practice estimates (CP_1_/CP_0_) by CP_ARB_. To avoid this problem, we directly used CP_1_ to calculate the crediting error for CAR1183. Using this method, we estimated that 100% of the project’s upfront credits were over-credited.

In light of this methodological nuance and the unusual situation of the addition of a new assessment area, we performed a complementary analysis to assess the robustness of our 100% over-crediting result. Instead of asking what would happen if the project baseline had been subjected to our alternative common practice estimate, we asked instead whether the project would have earned any upfront credits under the terms of the 2015 US Forest Projects protocol that would have been required had the project paperwork been filed a few weeks later.

From the project documentation, we know that 14.1% of the project’s basal area is made up of Pinyon/Juniper species. Under the 2015 protocol, the Central New Mexico Pinyon/Juniper assessment area has a common practice of 8.74 tCO_2_e per acre and the Central New Mexico “Mixed Conifer” assessment area has a common practice of 42.77 tCO_2_e per acre. While we do not know the true acreage that should be classified as Pinyon/Juniper, if we assume that 14.1% of the project acreage would have been assigned to the Pinyon/Juniper assessment area and the remainder to the Mixed Conifer assessment area, the project’s realized common practice would be 37.97 tCO_2_e per acre [8.74 * 0.141 + 42.77 * (1-0.141)].

Because the project’s initial carbon stocks were 35.61 tCO_2_e per acre, which is less than 37.97 tCO_2_e per acre, our separate calculation indicates that the project would not have been eligible for upfront carbon credits under the terms of the 2015 US Forest Offsets protocol. This result is independent of, and thus complementary to, our primary reclassification-based analysis, which produces the same result.

For our alternate analysis, it is important to note that the CARB protocol requires that project acreage be assigned to assessment areas on an ‘area-weighted’, as opposed to ‘basal-area weighted’ basis. However, because the project came in under the 2014 protocol rules, when only a single assessment area (Pinyon/Juniper) existed, no such area-weighted breakdown is provided in the project’s documentation. Instead, we make the assumption that species-level basal area serves as a reasonable proxy for project area. This assumption has biases that cut in both directions. On the one hand, basal area could under-predict the project area that is Pinyon/Juniper woodland because these Pinyon/Juniper crowns can be relatively well-spaced and therefore take up a greater share of land to produce a given share of basal area. On the other hand, we know that the Pinyon and Juniper species listed on the initial OPDR co-associate with the Mixed Conifer forest type strata, specifically Ponderosa pine, so less than 100% of the basal area of Pinyon and Juniper species would classified as Pinyon/Juniper woodland. Thus, in the absence of additional information, we believe using basal area as a proxy for total acreage is a reasonable assumption. In order for initial carbon stocks to exceed the project’s common practice number, which is required to award any “upfront” credits to the project, Pinyon/Juniper would need to account for 20% of the total project acreage.

### Spatial arbitrage patterns

For purposes of understanding the finer spatial variations in carbon stocks, we also worked with FIA data on a “per condition” basis. The approach described here was used to create the arbitrage potential map in Figure 4A, but not used elsewhere in our analysis.

For each supersection, we began by loading all FIA conditions for that supersection, as well as all bordering supersections. We then filtered the FIA data to meet the following criteria: (i) classified as accessible forestland (COND_STATUS_CD == 1); (ii) that were measured between 2001 and 2015; and (iii) fell on privately owned land. Using publicly reported (e.g., fuzzed and swapped) plot latitude and longitude, we assigned each condition a mean temperature and mean precipitation based on 30-year climate normals from PRISM (7). PRISM data were first regridded to a 4km Albers Equal Area Projection using area weighted resampling. Though reported FIA coordinates are approximate, the uncertainty in plot location (within ∼500 acres) is comparable to the 4km^2^ spatial resolution of the regridded PRISM data. To account for the difference in magnitude between precipitation and mean annual temperature, we transformed both quantities using a quantile transformer, which maps the cumulative distribution function of observed data to a uniform distribution. Each value is mapped (via its quantile) to the new distribution. This approach aids in the comparison of values measured on different scales (here, millimeters and degrees Celsius). Intuitively, a 10 mm change in precipitation is much less drastic than a 10 °C change in temperature. A 10 mm change in precipitation would hardly affect the reported quantile of an observation, whereas a difference of 10 °C would cause a large change in the reported quantile. Quantizing both measurements facilitates the subsequent analysis of identifying FIA plots in analogous “climate space.” Then, for each point in the 4km PRISM climate grid, we looked up the nearest *n* points in climate space, where *n* was set equal to 10% of all conditions (of any forest type) found in (i) the supersection and (ii) its bordering supersections. We then calculated mean standing live above ground carbon across those *n* conditions, taking into account per tree expansions factors (TPA_UNADJ) and condition proportion (CONDPROP_UNADJ). In addition to mean standing live aboveground carbon, we also calculated “relative standing aboveground carbon” by dividing each 4km estimate of mean carbon by the mean of all FIA plots falling within the supersection (e.g., excluding conditions in bordering supersections).

We reiterate that none of our analysis of crediting error (e.g. Figure 3) uses FIA data on a “per condition” basis in the manner described above. Rather, our primary analysis strictly follows the sampling and stratification rules of FIA survey design per the open-source rFIA package methods. “Per condition” data are only used for demonstrative purposes in Figure 4A to highlight the distinct biogeography of carbon stocks in the Southern Cascades supersection.

### Open source software and data

We performed all analyses using Python and, in the case of rFIA (5), the R programming language (8) in the Pangeo cloud environment (9). Our workflow used the following open source software packages: Pandas (10); Xarray (11); Matplotlib (12); NumPy (13); Seaborn (14); Jupyter (15); Scikit-learn (6). The source code to reproduce our analysis is available in (16, 17). Archival versions of this project’s data products are available in (3, 18).

## Glossary

|  |  |
| --- | --- |
| **Additionality** | When used to describe an offset project, the claim that a project's climate benefits were induced by the offset credit, i.e. that project scenario climate benefits are "in addition to" what would have happened in the baseline scenario or that the climate benefits would not exist without the offset project's activities.<br>When used to describe an offsets program, the claim that the climate benefits achieved by the program are equal to or greater than the number of credits awarded to participating projects.<br>Additionality is a critical requirement for compliance offsets in particular because offsets that are used in compliance contexts allow higher emissions that are premised on offsets' additional climate benefits. |
| **Air Resources Board Offset Credits (ARBOCs)** | The name for the offset credits issued by the California Air Resources Board. ARBOCs are eligible for compliance use in the state's cap-and-trade program. Each ARBOC is worth 1 metric tCO<sub>2</sub>e. |
| **Assessment Area** | A forest type that spans the full geographic extent of a supersection. Each supersection contains one or more assessment areas, each with a distinct estimate of common practice that is based on the average carbon stock for this forest type from USFS FIA data. |
| **Basal area** | The cross-sectional area of a tree at breast height. Often used to describe the total cross-sectional area of all trees on a plot. |
| **Baseline scenario** | A carbon offset has a project scenario and a baseline scenario. The baseline scenario describes the emissions outcomes that would happen counterfactually in the absence of an offset project, i.e. what would happen if the offset project is not pursued. By definition, an offset project's baseline scenario cannot be observed because it does not occur. |
| **California Air Resources Board (CARB)** | California's climate change regulator, which is responsible for the state's cap-and-trade and compliance carbon offsets program. CARB imported the core architecture of the US Forest Project protocol from an earlier Climate Action Reserve protocol in its 2011 and 2014 US Forest Projects protocols, then subsequently updated common practice numbers in coordination with USFS for its 2015 protocol revision. |
| **Climate Action Reserve (CAR)** | A nonprofit organization and carbon offset registry that developed the original forest offset program subsequently adopted and then revised by CARB. CAR created supersections and assessment areas, and developed the original common practice numbers adopted without change by the 2011 and 2014 CARB US Forest Projects protocols. |
| **Common practice** | The average carbon stock (tCO<sub>2</sub>/acre) in a given assessment area, as calculated across an entire supersection. |
| **Compliance offsets** | Carbon offsets that are fungible for compliance purposes in legally binding climate mitigation policies. Here, California has a cap-and-trade program that sets aggregate limits on greenhouse gas pollution. Because forest carbon offsets can be used by polluters to comply with the cap-and-trade program, forest carbon offsets are known as compliance offsets. Compliance offsets increase the total emissions allowed under climate mitigation policies premised on their claim to generate equivalent climate benefits elsewhere. |
| **CP<sub>ARB</sub>** | Common practice as reported by individual projects in the CARB program, expressed in tCO<sub>2</sub>/acre as a weighted average of all project assessment areas. |
| **CP<sub>0</sub>** | This study's re-calculation of common practice directly from FIA data, matching methods used by CARB and expressed in tCO<sub>2</sub>/acre. |
| **CP<sub>1</sub>** | This study's re-calculation of common practice using a classification model to match projects' actual species with comparable FIA plots in order to minimize ecological bias, expressed in tCO<sub>2</sub>/acre. |
| **Ecosection** | A geographic region defined by the USFS. CAR combined multiple individual ecosections together to form supersections, which serve as the geographic aggregations across which assessment areas are defined and common practice is calculated. |
| **Initial carbon stock (ICS)** | A measure of the standing live aboveground carbon stock in a given forest. Improved forest management (IFM) projects receive large upfront credits in their first reporting period (RP1) when their ICS exceeds the 100-year average carbon stock in their baseline scenario. |
| **Improved Forest Management (IFM)** | A kind of forest offset project that claims to increase average carbon stocks over time by changing the rotation or other management techniques affecting forest growth and harvest cycles. In California's US Forest Projects protocol, IFM credits receive "upfront" credit based on the difference between their initial standing carbon stock (as measured by site surveys) and the 100-year average of carbon stocks in their projected counterfactual baseline scenario. For IFM projects that have initial standing carbon stock above common practice, the 100-year average of carbon stocks in their baseline scenario must be equal to or greater to common practice. |
| **IFM-1** | A component of the "GHG boundary" for which greenhouse gas emission reductions can be credited, representing onsite standing live tree carbon, both above and belowground, in either the baseline or project scenario. Reported in an offset project's OPDR. |
| **IFM-3** | A component of the "GHG boundary" for which greenhouse gas emission reductions can be credited, representing the amount of onsite standing dead tree carbon, in either the baseline or project scenario. Reported in an offset project's OPDR. |
| **IFM-7** | A component of the “GHG boundary” for which greenhouse gas emission reductions can be credited, representing the amount of carbon stored within “in-use” wood products, in either the baseline or project scenario. Reported in an offset project’s OPDR. |
| **IFM-8** | A component of the “GHG boundary” for which greenhouse gas emission reductions can be credited, representing the amount of carbon stored within “land-filled” wood products, in either the baseline or project scenario. Reported in an offset project’s OPDR. |
| **Forest Inventory and Analysis (FIA)** | A comprehensive dataset describing forests and forest ecology in the United States, collected and maintained by the US Forest Service. |
| **Non-additional** | When a project or program fails to achieve the additionality standard. |
| **Offset credit** | In the context of California’s program, an offset credit represents 1 tCO<sub>2</sub>e using 100-year IPCC AR4 global warming potentials. In California’s program, offset credits are called Air Resources Board Offset Credits (ARBOCs). |
| **Offset project data reports (OPDRs)** | A standardized public reporting document required for each offset project and filed with one of the private carbon offset registries that helps CARB implement its compliance offset program. OPDRs contain critical information about offset projects, including species-level data, baseline, and project scenario information, and serve as the basis for the calculations a project developer makes to claim credits from CARB. Using a critical subset of these data (IFM-1, IFM-3, IFM-7, and IFM-8) we are able to re-calculate the number of offset credits that should be issued to any publicly listed offset project with an OPDR. |
| **Over-crediting** | The outcome in which a project is awarded more credits than the climate benefits it can rightly claim. In this study, we report over-crediting when our re-estimate of common practice leads to a higher number than what a project uses to earn credits under the US Forest Projects protocol. |
| **Project scenario** | A carbon offset has a project scenario and a baseline scenario. The project scenario is the scenario that describes the emissions outcomes when an offset project is implemented, i.e. what a project claims will happen in reality if pursued. |
| **RP1** | Reporting Period 1, the first reporting period for offset projects’ documentation. All upfront credits are awarded in RP1, along with the first tranche of annual credits that reflect forest growth. |
| **RP2** | Reporting Period 2, the second reporting period for offsets’ project documentation. The second tranche of annual credits that reflect forest growth are awarded in RP2. Used here to help verify crediting calculations. |
| **Standardized approach to carbon offsets** | A paradigm for offset program regulation. Earlier offset programs used project-level calculations and bespoke methods that regulators approved |
|  | on a case-by-case basis. Standardized offset programs have common rules that establish what kinds of projects are eligible, how project baselines scenarios are determined, and how to calculate the number of offset credits an eligible project should earn. |
| **Supersection** | The geographic unit of analysis in California's forest offsets protocol. CAR originally created supersections in the continental United States by combining together multiple ecosections to form supersections. Each supersection has one or more assessment areas, each with a distinct estimate of common practice. CARB adopted CAR's supersections for use in its compliance offsets program, adding Alaska in its 2015 US Forest Projects protocol. |
| **US Forest Projects protocol** | The California Air Resources Board's forest offset protocol, which sets out standardized rules for project eligibility, baseline, and crediting calculations. CARB has adopted three versions of the protocol: the first in 2011, the second in 2014, and the third and current version in 2015. This protocol produces over 80% of the offset credits in California's cap-and-trade program. All versions have been open to projects anywhere in the continental United States; as of the 2015 version, projects in southern coastal Alaska became eligible as well. |
| **US Forest Service (USFS)** | A branch of the US Department of Agriculture that is responsible for collecting and maintaining the FIA data used here (among other matters). |
| **Under-crediting** | The outcome in which a project is awarded fewer credits than the climate benefits it can rightly claim. In this study, we report under-crediting when our re-estimate of common practice leads to a lower number than what a project uses to earn credits under the US Forest Projects protocol. |
| **Upfront credits** | The credits received by an improved forest management (IFM) project in its first reporting period (RP1). In RP1, most OPDRs do not distinguish between (1) annual forest growth and (2) the much larger number of credits awarded to IFM projects with initial carbon stocks above 100-year average carbon stocks in projects' baselines, so we report both components as upfront credits. |
| **Voluntary offsets** | Carbon offsets that are bought and sold for voluntary, typically private purposes, such as an individual or company wanting to claim carbon neutrality. California's offsets program is a compliance offsets program, not a voluntary program. |

## Appendix 1: Verification of crediting calculations

Here, we describe a project-by-project description of projects where we identified a discrepancy between the number of credits a project’s documentation claims in its official reporting (ARBOC_Reported_), the number of credits th California Air Resources Board issues to the project (ARBOC_Issuance_), and our independent effort to recalculate the appropriate number of credits from projects documentation (ARBOC_Calculated_).

The discrepancies can be categorized in two groups. The first group includes projects for which the number of offset credits reported by offset projects is not equal to what the regulator issued (ARBOC_Reported_ ≠ ARBOC_Issuance_), and the second group includes projects for which the number of offset credits reported by offset projects is not equal to what our independent calculations (ARBOC_Reported_ ≠ ARBOC_Calculated_). We address each in turn.

### Reported not equal to issuance

We start with cases where ARBOC_Reported_ is not equal to ARBOC_Issuance_, further subdividing these cases into four sub-groupings. For each sub-grouping, we list each instance of a discrepancy.

#### Unexplained — potential over-crediting

|  |  |
| --- | --- |
| **CAR1175** | Difference of 30 ARBOCs between Reported and Issuance. RP1 has two verification statements — one for 3,824,257 and another for 3,824,227. The annual OPDR for RP2 also records OPDRreported of 3,824,227 ARBOCs for RP1. It is possible that CARB over-issued 30 ARBOCs. It does not appear 30 ARBOCs were deducted from a later reporting period. Both G. Badgley and B. Haya calculate a 30 ARBOC difference. |

#### Unexplained — possible out–of–date documents

In all the instances listed below, we’ve tried to triangulate what the project owner/developer formally requested from CARB. It is our impression that these five projects have updated OPDRs that have not been posted to the registries.

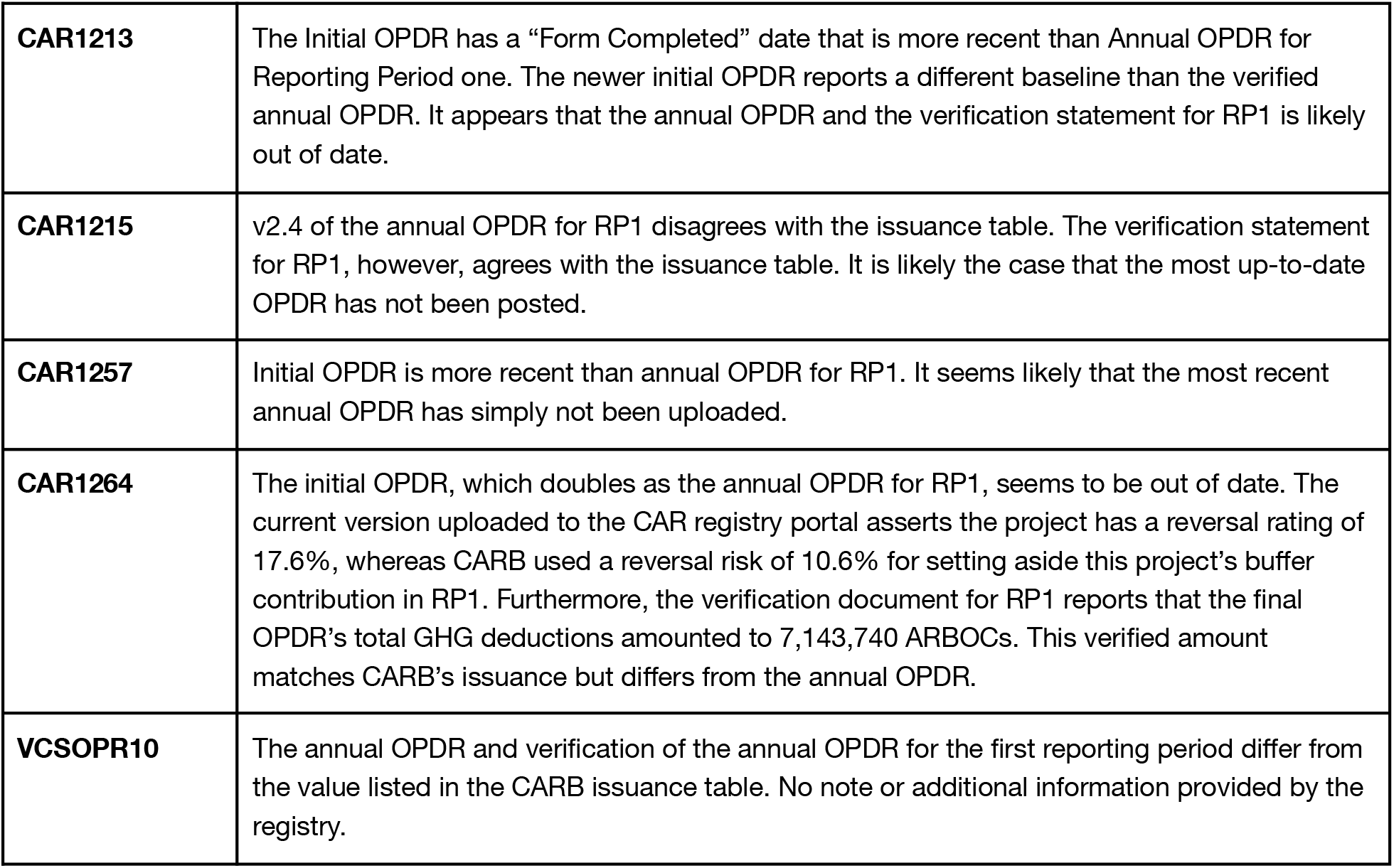

#### Correctable Errors

Two discrepancies arise from projects that have “Correctable Error” notes including in the project documents listed at the registry. These notes indicate that the regulator has taken an action to modify the number of credits reported by the project, but without additional explanation.

|  |  |
| --- | --- |
| **CAR1103** | On 11/29/2016, CAR issued a Project Note that states that "During [CARB's] regulatory review, [CARB] identified a correctable error." The note specifies that 270,943 ARBOCs were issued, a number that matches the CARB issuance table. We could not identify a copy of the updated/corrected OPDR. |
| **CAR1208** | On 5/14/20, CAR issued a Project Note that states that "During [CARB's] regulatory review, [CARB] identified a correctable error." The note specifies that 501,850 ARBOCs were issued, a number that matches the CARB issuance table. We could not identify a copy of the updated/corrected OPDR. |

### Reported not equal to calculated

We now move to cases where ARBOC_Reported_ is not equal to ARBOC_Calculated_, again subdividing discrepancies into relevant sub-groupings.

#### Rounding confidence deduction

Projects report a confidence deduction to adjust for uncertainty estimates of onsite carbon stocks. We identified three projects where the confidence deduction has been rounded, causing differences between ARBOC_Reported_ and ARBOC_Calculated_.

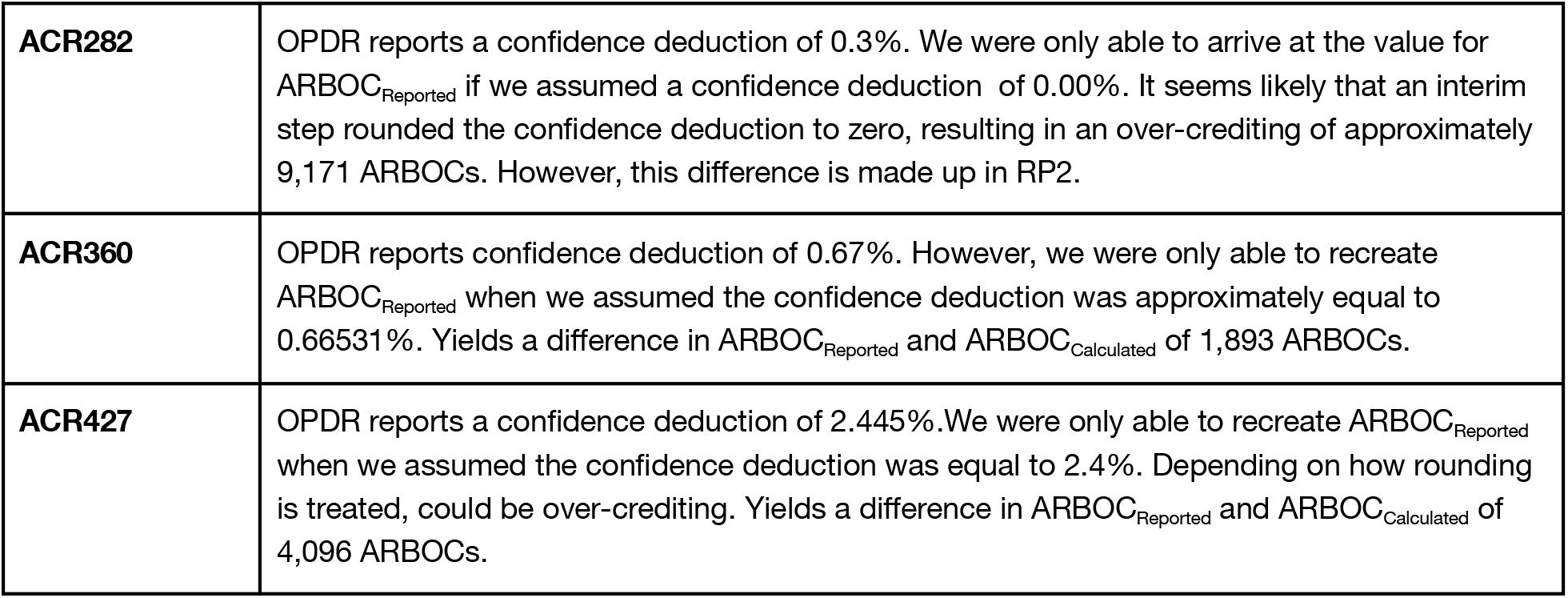

#### Harvest

We struggled to exactly replicate the crediting calculation in the following three cases that share two attributes: (i) significant harvesting in the project scenario combined with (ii) first reporting periods (RP1) of longer than one year. The longer reporting periods pose trouble because they introduce the possibility that the baseline wood products components (IFM-7 and IFM-8) need to be prorated. Prorating adds an extra difficulty because some project OPDRs reported the prorated values of IFM-7 and IFM-8, while others report the annual values, but appear to use prorated values in their underlying ARBOC calculations. Given the possibility of other reporting errors, this extra “degree of freedom” complicates reproducing ARBOC_Reported_. Getting baseline wood products correct is important because of how the protocol treats leakage when wood products generated in the harvest scenario exceed wood products generated in the counterfactual baseline scenario. Getting leakage calculations correct is further complicated by potential differences in the quality/composition of wood products in the product vs baseline scenario. Combined together, these discrepancies make it difficult to recreate the issuance calculations with a high degree of confidence.

In all cases we get ARBOC_Calculated_ exceeding ARBOC_Reported_, meaning there is little likelihood of over-issuance. Ideally, the need to prorate IFM-7 and IFM-8 should not be required to recreate issuance calculations, but current CARB reporting requirements do not appear to strictly enforce the time horizon over which IFM-7 and IMF-8 are reported in annual OPDRs.

|  |  |
| --- | --- |
| **ACR247** | Large harvest component. Unable to replicate issuance after attempting to make reasonable pro-rating assumptions. We estimate $ARBOC_{Calculated}$ as exceeding $ARBOC_{Reported}$ by 12,947 ARBOCs. Discrepancy likely has something to do with how the project prorated harvest in baseline and potentially how the project calculated secondary effects. RP1 duration of ~2 years. |
| **CAR1217** | Large harvest component. We estimate $ARBOC_{Calculated}$ exceeding $ARBOC_{Reported}$ by 1,047 ARBOCs. RP1 duration of ~2 years. |
| **ACR276** | Large harvest component. We estimate $ARBOC_{Calculated}$ exceeding $ARBOC_{Reported}$ by 3,298 ARBOCs. RP1 duration of > 1 year. |

#### Errors under 25 ARBOCs that might explained by confidence deduction rounding

All these projects have smaller differences in ARBOC_Reported_ as compared to ARBOC_Calculated_. However, all these projects also have a confidence deduction greater than 0. Therefore, we cannot rule out that rounding of the confidence deduction is the source of the difference. Differences reported below have been rounded.

|  |  |
| --- | --- |
| **CAR1094** | Off by 3. |
| **CAR1204** | Off by 14. |
| **ACR256** | Off by 21. |
| **ACR257** | Off by 2. |

#### Errors under 25 ARBOCs that cannot be explained by confidence deduction rounding

This project is off by precisely two. Likely a data entry issue somewhere in the OPDR.

|  |  |
| --- | --- |
| **CAR1032** | Off by 2; whole value so likely not rounding. |

#### De minimis errors (< 2 ARBOC) that can be explained by leakage/CD rounding

All these projects have *even smaller* differences in ARBOC_Reported_ as compared to ARBOC_Calculated_. However, all these projects also have a confidence deduction greater than 0. Therefore, we cannot rule out rounding of the confidence dedication as the source of the difference.

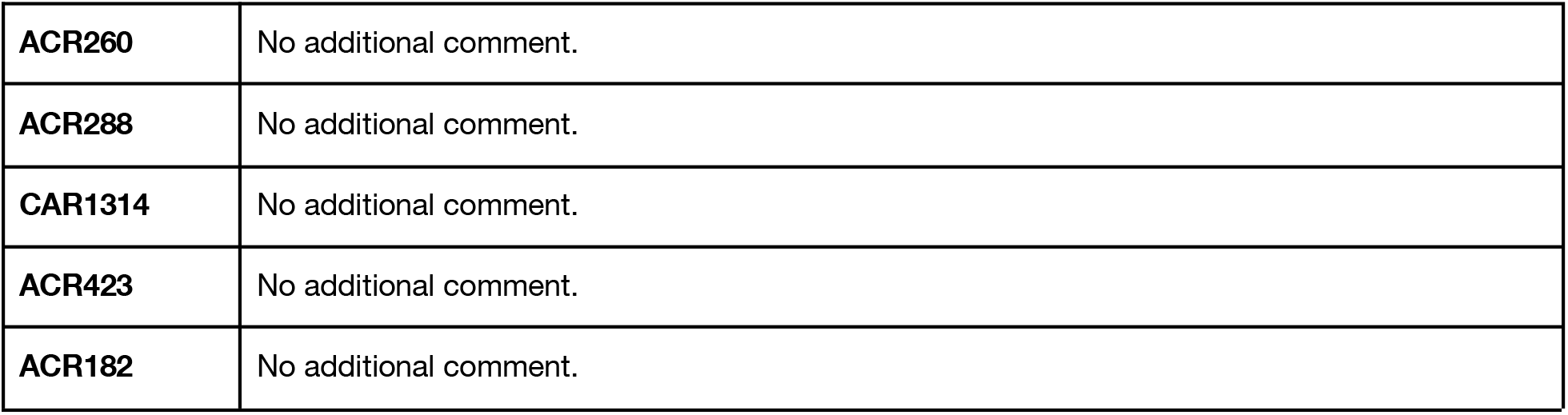

#### Errors of less than or equal to two, not explained by confidence deduction

These are projects where the confidence deduction of the first reporting period is zero. That means rounding cannot fully explain the difference. It’s still possible that intermediate rounding of leakage on wood products could partially explain these differences.

|  |  |
| --- | --- |
| **CAR1066** | No additional comment. |
| **ACR393** | Off by exactly 1. |

## Appendix 2: Classification labels

Here we provide species composition and output of our classifier (described above in “Classification algorithm”) for all of the 65 projects included in the crediting error analysis reported in Figure 3. For brevity, we exclude listings in the “Project species” and “Forest type classification” columns that fall below 10% from this table; however, all digitized listings are used in the underlying analysis and available as part of our public data.

| Project | Supersection | Assessment Area | Project species (fractional basal area) | Forest type classification probabilities |
| --- | --- | --- | --- | --- |
| CAR1205 | 2 | 999 | Chestnut oak : 15.5%<br>Yellow-poplar : 14.0% | Yellow-poplar / white oak / northern red oak : 19.8%<br>White oak / red oak / hickory : 34.8%<br>Chestnut oak / black oak / scarlet oak : 38.1% |
| CAR1205 | 76 | 999 | Chestnut oak : 15.5%<br>Yellow-poplar : 14.0% | Yellow-poplar / white oak / northern red oak : 18.8%<br>Chestnut oak / black oak / scarlet oak : 34.8%<br>White oak / red oak / hickory : 39.7% |
| VCSOPR10 | 15 | 999 | Black walnut : 15.2%<br>Shortleaf pine : 15.6%<br>White oak : 29.2% | Shortleaf pine / oak : 12.5%<br>White oak : 16.9%<br>White oak / red oak / hickory : 68.2% |
| ACR192 | 4 | 999 | Longleaf pine : 32.9%<br>Loblolly pine : 19.3%<br>Laurel oak : 18.6% | Longleaf pine / oak : 15.1%<br>Loblolly pine : 15.8%<br>Longleaf pine : 63.7% |
| ACR247 | 1 | 1 | Balsam fir : 38.2%<br>Red maple : 17.6%<br>Yellow birch : 14.3% | Sugar maple / beech / yellow birch : 83.0% |
| ACR247 | 1 | 2 | Red maple : 17.7%<br>Sugar maple : 30.8%<br>Yellow birch : 13.8%<br>Beech : 14.5% | Sugar maple / beech / yellow birch : 99.0% |
| ACR262 | 55 | 221 | Douglas fir : 35.0%<br>Tanoak : 29.0% | Douglas fir : 22.9%<br>Tanoak : 64.6% |
| ACR262 | 79 | 297 | Douglas fir : 35.0% | Douglas fir : 18.1% |
|  |  |  | Tanoak : 29.0% | Tanoak : 72.8% |
| ACR257 | 32 | 113 | Loblolly pine : 64.8% | Loblolly pine : 92.2% |
| ACR280 | 2 | 999 | Chestnut oak : 11.5%<br>Red maple : 10.8%<br>Yellow-poplar : 14.5% | Chestnut oak / black oak / scarlet oak : 25.4%<br>White oak / red oak / hickory : 28.7%<br>Yellow-poplar / white oak / northern red oak : 31.4% |
| CAR1190 | 55 | 221 | Tanoak : 23.0%<br>Redwood : 49.0%<br>Douglas fir : 24.0% | Tanoak : 26.6%<br>Douglas fir : 28.5%<br>Redwood : 39.7% |
| CAR1208 | 2 | 3 | Yellow-poplar : 27.2%<br>Eastern hemlock : 24.4%<br>Virginia pine : 17.1% | Virginia pine / southern red oak : 13.5%<br>Yellow-poplar / white oak / northern red oak : 52.5% |
| CAR1208 | 2 | 4 | Yellow-poplar : 21.6%<br>Sycamore : 10.6% | Cherry / white ash / yellow-poplar : 10.4%<br>White oak / red oak / hickory : 20.8%<br>Yellow-poplar / white oak / northern red oak : 46.5% |
| CAR1208 | 2 | 7 | Sugar maple : 17.6%<br>Yellow-poplar : 14.4% | Sugar maple / beech / yellow birch : 11.3%<br>Yellow-poplar / white oak / northern red oak : 33.0%<br>White oak / red oak / hickory : 35.5% |
| CAR1208 | 2 | 8 | Chestnut oak : 14.8%<br>White oak : 10.9%<br>Black oak : 10.6% | White oak / red oak / hickory : 42.5%<br>Chestnut oak / black oak / scarlet oak : 47.5% |
| CAR1208 | 2 | 5 | White oak : 24.2%<br>Beech : 22.4%<br>Yellow-poplar : 12.3%<br>Red maple : 11.0% | Yellow-poplar / white oak / northern red oak : 35.4%<br>White oak / red oak / hickory : 51.0% |
| CAR1208 | 24 | 86 | Yellow-poplar : 27.2%<br>Eastern hemlock : 24.4%<br>Virginia pine : 17.1% | Yellow-poplar / white oak / northern red oak : 58.4% |
| CAR1208 | 24 | 87 | Yellow-poplar : 21.6%<br>Sycamore : 10.6% | White oak / red oak / hickory : 23.1%<br>Yellow-poplar / white oak / northern red oak : 43.1% |
| CAR1208 | 24 | 89 | Sugar maple : 17.6%<br>Yellow-poplar : 14.4% | Yellow-poplar / white oak / northern red oak : 32.9%<br>White oak / red oak / hickory : 40.0% |
| CAR1208 | 24 | 90 | Chestnut oak : 14.8%<br>White oak : 10.9%<br>Black oak : 10.6% | White oak / red oak / hickory : 44.7%<br>Chestnut oak / black oak / scarlet oak : 46.0% |
| CAR1208 | 24 | 92 | White oak : 24.2%<br>Beech : 22.4%<br>Yellow-poplar : 12.3%<br>Red maple : 11.0% | Yellow-poplar / white oak / northern red oak : 30.0%<br>White oak / red oak / hickory : 57.6% |
| CAR1264 | 287 | 287 | Sitka spruce : 40.9%<br>Western hemlock : 29.2%<br>Mountain hemlock : 23.1% | Mountain hemlock : 19.5%<br>Sitka spruce : 34.3%<br>Western hemlock : 46.1% |
| ACR393 | 39 | 151 | Northern white-cedar : 54.0% | Northern white-cedar : 85.9% |
| ACR393 | 39 | 152 | Eastern hemlock : 15.0%<br>Red maple : 21.0%<br>Sugar maple : 31.0%<br>Yellow birch : 11.0% | Sugar maple / beech / yellow birch : 99.2% |
| ACR393 | 39 | 153 | Balsam fir : 12.0%<br>Eastern hemlock : 18.0%<br>Eastern whitepine : 12.0%<br>Northern white-cedar : 20.0%<br>Red maple : 23.0% | Northern white-cedar : 15.1%<br>Sugar maple / beech / yellow birch : 48.4% |
| ACR423 | 58 | 229 | Douglas fir : 71.0%<br>Western hemlock : 13.0% | Douglas fir : 95.2% |
| CAR1314 | 60 | 231 | Ponderosa pine : 57.0% | Ponderosa pine : 43.9% |
|  |  |  | Douglas fir : 57.0% | Douglas fir : 55.6% |
| CAR1314 | 60 | 232 | Ponderosa pine : 66.0%<br>Douglas fir : 30.0% | Douglas fir : 11.8%<br>Ponderosa pine : 87.3% |
| CAR1314 | 22 | 82 | Ponderosa pine : 63.0% | Ponderosa pine : 94.1% |
| ACR425 | 286 | 286 | Western hemlock : 79.0%<br>White spruce : 14.0% | Western hemlock : 99.0% |
| ACR427 | 94 | 347 | Balsam fir : 17.2%<br>Black spruce : 16.0%<br>Red spruce : 11.5%<br>Northern white-cedar : 20.0% | Black spruce : 10.0%<br>Balsam fir : 71.0% |
| ACR458 | 286 | 286 | Western hemlock : 45.0%<br>Western red cedar : 30.0%<br>Sitka spruce : 11.0% | Western redcedar : 27.8%<br>Western hemlock : 72.2% |
| CAR1180 | 55 | 221 | Douglas fir : 34.0%<br>Redwood : 13.0%<br>Tanoak : 24.0% | Douglas fir : 33.0%<br>Tanoak : 55.5% |
| ACR249 | 2 | 999 | Chestnut oak : 12.2%<br>Yellow-poplar : 16.0% | White oak / red oak / hickory : 26.3%<br>Chestnut oak / black oak / scarlet oak : 27.9%<br>Yellow-poplar / white oak / northern red oak : 36.7% |
| CAR1204 | 42 | 169 | Red maple : 31.4%<br>Sugar maple : 10.5% | Sugar maple / beech / yellow birch : 98.3% |
| CAR1204 | 42 | 171 | Aspen : 32.3%<br>Red maple : 13.2%<br>Balsam fir : 11.7%<br>Red spruce : 10.1% | Sugar maple / beech / yellow birch : 30.2%<br>Aspen : 69.8% |
| CAR1204 | 42 | 172 | Red spruce : 24.8%<br>Northern white-cedar : 19.1%<br>Red maple : 15.1%<br>Balsam fir : 14.2% | Sugar maple / beech / yellow birch : 10.9%<br>Balsam fir : 10.9%<br>Red spruce / balsam fir : 22.9%<br>Red spruce : 49.0% |
| CAR1204 | 94 | 347 | Red spruce : 24.8%<br>Northern white-cedar : 19.1%<br>Red maple : 15.1%<br>Balsam fir : 14.2% | Balsam fir : 11.0%<br>Sugar maple / beech / yellow birch : 11.8%<br>Red spruce / balsam fir : 21.2%<br>Red spruce : 49.5% |
| CAR1204 | 94 | 348 | Red maple : 31.4%<br>Sugar maple : 10.5% | Sugar maple / beech / yellow birch : 97.8% |
| ACR200 | 55 | 221 | Tanoak : 36.0%<br>Douglas fir : 32.0%<br>Redwood : 21.0% | Douglas fir : 21.6%<br>Tanoak : 75.4% |
| ACR256 | 24 | 999 | Chestnut oak : 29.1% | White oak / red oak / hickory : 17.4%<br>Chestnut oak : 20.7%<br>Chestnut oak / black oak / scarlet oak : 56.2% |
| ACR256 | 69 | 999 | Chestnut oak : 29.1% | White oak / red oak / hickory : 20.3%<br>Chestnut oak : 23.7%<br>Chestnut oak / black oak / scarlet oak : 46.5% |
| CAR1013 | 55 | 999 | Tanoak : 30.9%<br>Douglas fir : 27.8%<br>Redwood : 16.7% | Douglas fir : 20.3%<br>Tanoak : 74.6% |
| CAR1213 | 1 | 1 | Red spruce : 40.9%<br>Balsam fir : 16.3%<br>Blasck cherry : 11.2% | Sugar maple / beech / yellow birch : 10.8%<br>Red spruce : 14.3%<br>Red spruce / balsam fir : 68.3% |
| CAR1213 | 1 | 2 | Red maple : 12.2%<br>Beech : 31.4%<br>Sugar maple : 13.3%<br>Yellow birch : 14.9% | Sugar maple / beech / yellow birch : 100.0% |
| CAR1213 | 86 | 318 | Red spruce : 40.9%<br>Balsam fir : 16.3%<br>Blasck cherry : 11.2% | Sugar maple / beech / yellow birch : 11.4%<br>Red spruce : 13.2%<br>Red spruce / balsam fir : 69.6% |
| CAR1213 | 86 | 319 | Red maple : 12.2%<br>Beech : 31.4%<br>Sugar maple : 13.3%<br>Yellow birch : 14.9% | Sugar maple / beech / yellow birch : 100.0% |
| ACR199 | 94 | 348 | Sugar maple : 39.5%<br>Yellow birch : 25.9% | Sugar maple / beech / yellow birch : 100.0% |
| ACR199 | 94 | 347 | Balsam fir : 38.1%<br>Red spruce : 17.7%<br>Yellow birch : 17.7% | Balsam fir : 12.9%<br>Red spruce / balsam fir : 26.7%<br>Sugar maple / beech / yellow birch : 47.0% |
| ACR182 | 55 | 221 | Douglas fir : 34.2%<br>Redwood : 15.8%<br>Pacific madrone : 11.6%<br>Tanoak : 25.8% | Douglas fir : 27.7%<br>Tanoak : 59.2% |
| ACR279 | 2 | 999 | Chestnut oak : 11.6%<br>Red maple : 19.0%<br>Yellow-poplar : 18.4% | White oak / red oak / hickory : 18.4%<br>Chestnut oak / black oak / scarlet oak : 25.1%<br>Yellow-poplar / white oak / northern red oak : 31.3% |
| ACR267 | 2 | 999 | Chestnut oak : 12.8%<br>Yellow-poplar : 17.7% | White oak / red oak / hickory : 22.7%<br>Chestnut oak / black oak / scarlet oak : 26.5%<br>Yellow-poplar / white oak / northern red oak : 36.4% |
| CAR1183 | 18 | 71 | Douglas fir : 37.1%<br>Ponderosa pine : 22.9%<br>White fir : 12.8% | Ponderosa pine : 28.5%<br>Douglas fir : 68.4% |
| ACR202 | 38 | 999 | Eastern hemlock : 11.9%<br>Northern white-cedar : 16.5%<br>Sugar maple : 34.5% | Hard maple / basswood : 18.1%<br>Sugar maple / beech / yellow birch : 81.9% |
| ACR202 | 35 | 999 | Eastern hemlock : 11.9%<br>Northern white-cedar : 16.5%<br>Sugar maple : 34.5% | Hard maple / basswood : 12.8%<br>Sugar maple / beech / yellow birch : 87.2% |
| ACR361 | 287 | 287 | Sitka spruce : 94.9% | Sitka spruce : 98.0% |
| CAR1197 | 1 | 1 | Eastern hemlock : 23.2%<br>Red spruce : 20.2%<br>Red maple : 11.1%<br>Balsam fir : 11.0% | Red spruce : 12.0%<br>Eastern hemlock : 18.7%<br>Sugar maple / beech / yellow birch : 58.3% |
| CAR1197 | 1 | 2 | Beech : 33.7%<br>Sugar maple : 28.2%<br>Yellow birch : 10.8% | Sugar maple / beech / yellow birch : 99.8% |
| CAR1197 | 86 | 318 | Eastern hemlock : 23.2%<br>Red spruce : 20.2%<br>Red maple : 11.1%<br>Balsam fir : 11.0% | Red spruce : 15.0%<br>Eastern hemlock : 21.9%<br>Sugar maple / beech / yellow birch : 53.6% |
| CAR1197 | 86 | 319 | Beech : 33.7%<br>Sugar maple : 28.2%<br>Yellow birch : 10.8% | Sugar maple / beech / yellow birch : 99.8% |
| CAR973 | 37 | 999 | Eastern hemlock : 10.7%<br>Red maple : 25.9%<br>Sugar maple : 28.0% | Sugar maple / beech / yellow birch : 95.6% |
| CAR973 | 39 | 999 | Eastern hemlock : 10.7%<br>Red maple : 25.9%<br>Sugar maple : 28.0% | Sugar maple / beech / yellow birch : 98.3% |
| CAR1066 | 79 | 297 | White fir : 71.8%<br>Red cedar : 13.9% | California mixed conifer : 18.5%<br>White fir : 78.1% |
| CAR1041 | 79 | 297 | Douglas fir : 58.7% | California mixed conifer : 16.0%<br>Douglas fir : 78.0% |
| CAR1104 | 79 | 297 | Douglas fir : 35.6%<br>Tanoak : 28.5%<br>Pacific madrone : 15.0% | Douglas fir : 17.5%<br>Tanoak : 69.9% |
| ACR173 | 79 | 297 | Douglas fir : 56.4%<br>Pacific madrone : 12.1%<br>California black oak : 10.1% | California mixed conifer : 21.5%<br>Douglas fir : 67.2% |
| CAR1191 | 55 | 221 | Douglas fir : 29.4%<br>Redwood : 20.0% | Douglas fir : 11.9%<br>Tanoak : 86.8% |
|  |  |  | Tanoak : 43.0% |  |
| ACR210 | 39 | 151 | Eastern hemlock : 19.4%<br>Northern white-cedar : 46.3% | Northern white-cedar : 79.1% |
| ACR210 | 39 | 152 | Sugar maple : 23.5%<br>Red maple : 12.6%<br>Eastern hemlock : 42.8% | Sugar maple / beech / yellow birch : 89.4% |
| ACR210 | 39 | 153 | Northern red oak : 12.1%<br>Eastern whitepine : 43.6%<br>Red pine : 15.2% | Eastern white pine / northern red oak / white ash : 29.6%<br>Eastern white pine : 64.5% |
| ACR210 | 39 | 154 | Quaking aspen : 10.1%<br>Red maple : 13.4%<br>Eastern hophornbeam : 12.4%<br>Eastern whitepine : 14.0%<br>Aspen : 20.0% | Sugar maple / beech / yellow birch : 12.3%<br>Aspen : 82.1% |
| ACR276 | 2 | 999 | Chestnut oak : 11.0%<br>Red maple : 13.0% | Yellow-poplar / white oak / northern red oak : 21.7%<br>Chestnut oak / black oak / scarlet oak : 25.8%<br>White oak / red oak / hickory : 38.5% |
| ACR276 | 76 | 999 | Chestnut oak : 11.0%<br>Red maple : 13.0% | Yellow-poplar / white oak / northern red oak : 20.6%<br>Chestnut oak / black oak / scarlet oak : 21.1%<br>White oak / red oak / hickory : 44.1% |
| ACR189 | 79 | 297 | Douglas fir : 26.0%<br>Tanoak : 49.0% | Tanoak : 95.9% |
| ACR303 | 95 | 999 | Douglas fir : 11.9%<br>Ponderosa pine : 43.0% | Ponderosa pine : 95.5% |
| ACR303 | 88 | 999 | Douglas fir : 11.9% | Ponderosa pine : 85.2% |
|  |  |  | Ponderosa pine :<br>43.0% |  |
| CAR1173 | 24 | 999 | Sugar maple : 20.1%<br>Yellow-poplar : 18.7%<br>Chestnut oak : 12.5%<br>White oak : 11.6% | Chestnut oak / black oak / scarlet oak : 10.9%<br>Yellow-poplar / white oak / northern red oak : 37.1%<br>White oak / red oak / hickory : 37.8% |
| CAR1173 | 76 | 999 | Sugar maple : 20.1%<br>Yellow-poplar : 18.7%<br>Chestnut oak : 12.5%<br>White oak : 11.6% | Yellow-poplar / white oak / northern red oak : 34.5%<br>White oak / red oak / hickory : 39.7% |
| CAR1046 | 79 | 999 | Douglas fir : 35.7%<br>White fir : 22.6%<br>Tanoak : 13.9% | California mixed conifer : 37.6%<br>Douglas fir : 49.5% |
| CAR1209 | 35 | 999 | American basswood : 19.3%<br>Sugar maple : 56.1% | Sugar maple / beech / yellow birch : 16.3%<br>Hard maple / basswood : 83.5% |
| CAR1209 | 38 | 999 | American basswood : 19.3%<br>Sugar maple : 56.1% | Sugar maple / beech / yellow birch : 23.5%<br>Hard maple / basswood : 76.4% |
| CAR1257 | 2 | 3 | Chestnut oak : 31.2%<br>Northern red oak : 20.6%<br>Red maple : 20.0% | White oak / red oak / hickory : 18.0%<br>Chestnut oak : 21.9%<br>Chestnut oak / black oak / scarlet oak : 50.1% |
| CAR1257 | 2 | 4 | Chestnut oak : 15.8%<br>Yellow-poplar : 15.6%<br>Red maple : 14.9% | White oak / red oak / hickory : 20.7%<br>Yellow-poplar / white oak / northern red oak : 28.5%<br>Chestnut oak / black oak / scarlet oak : 39.1% |
| CAR1257 | 2 | 7 | Sugar maple : 22.9%<br>Yellow-poplar : 16.0% | Hard maple / basswood : 13.3%<br>Sugar maple / beech / yellow birch : 15.4%<br>Yellow-poplar / white oak / northern red oak : 25.7%<br>White oak / red oak / hickory : 29.8% |
| CAR1257 | 2 | 8 | Chestnut oak : 21.2%<br>Northern red oak : 12.9%<br>Yellow-poplar : 11.5%<br>Pignut hickory : 11.3%<br>Red maple : 10.2% | Yellow-poplar / white oak / northern red oak : 13.6%<br>White oak / red oak / hickory : 32.8%<br>Chestnut oak / black oak / scarlet oak : 45.4% |
| CAR1257 | 76 | 292 | Sugar maple : 22.9%<br>Yellow-poplar : 16.0% | Hard maple / basswood : 12.4%<br>Sugar maple / beech / yellow birch : 16.9%<br>Yellow-poplar / white oak / northern red oak : 24.9%<br>White oak / red oak / hickory : 30.4% |
| CAR1257 | 76 | 291 | Chestnut oak : 21.2%<br>Northern red oak : 12.9%<br>Yellow-poplar : 11.5%<br>Pignut hickory : 11.3%<br>Red maple : 10.2% | Yellow-poplar / white oak / northern red oak : 14.0%<br>White oak / red oak / hickory : 36.2%<br>Chestnut oak / black oak / scarlet oak : 41.9% |
| ACR284 | 1 | 1 | Eastern hemlock : 43.5%<br>Eastern whitepine : 22.3%<br>Red maple : 12.0% | Sugar maple / beech / yellow birch : 36.3%<br>Eastern white pine / eastern hemlock: 46.1% |
| ACR284 | 1 | 2 | Sugar maple : 20.9%<br>Red maple : 16.8%<br>Beech : 11.8%<br>Eastern whitepine : 10.3%<br>White ash : 10.3% | Sugar maple / beech / yellow birch : 88.8% |
| ACR284 | 41 | 162 | Red maple : 24.3%<br>Northern red oak : 20.3%<br>Eastern whitepine : 17.2%<br>Eastern hemlock : 16.5% | White oak / red oak / hickory : 20.3%<br>Sugar maple / beech / yellow birch : 58.3% |
| ACR284 | 41 | 163 | Eastern hemlock : 43.5%<br>Eastern whitepine : 22.3%<br>Red maple : 12.0% | Eastern white pine / eastern hemlock : 37.6%<br>Sugar maple / beech / yellow birch : 42.3% |
| ACR284 | 41 | 164 | Sugar maple : 20.9%<br>Red maple : 16.8%<br>Beech : 11.8%<br>Eastern whitepine : 10.3%<br>White ash : 10.3% | Sugar maple / beech / yellow birch : 87.3% |
| CAR1297 | 44 | 999 | Douglas fir : 75.7% | Douglas fir : 96.6% |
| CAR1215 | 2 | 3 | Eastern hemlock : 21.4%<br>Red maple : 17.6%<br>Scarlet oak : 10.1% | Sugar maple / beech / yellow birch : 19.9%<br>White oak / red oak / hickory : 20.7%<br>Chestnut oak / black oak / scarlet oak : 21.7%<br>Yellow-poplar / white oak / northern red oak : 25.3% |
| CAR1215 | 2 | 4 | Yellow-poplar : 19.5%<br>Red maple : 16.1% | Cherry / white ash / yellow-poplar : 12.2%<br>Chestnut oak / black oak / scarlet oak : 16.4%<br>White oak / red oak / hickory : 17.8%<br>Yellow-poplar / white oak / northern red oak : 36.1% |
| CAR1215 | 2 | 7 | Yellow-poplar : 17.6%<br>Sugar maple : 15.8%<br>Red maple : 10.7% | White oak / red oak / hickory : 25.5%<br>Yellow-poplar / white oak / northern red oak : 38.5% |
| CAR1215 | 2 | 8 | Chestnut oak : 33.0%<br>Red maple : 19.2% | Chestnut oak : 31.5%<br>Chestnut oak / black oak / scarlet oak : 58.0% |
| CAR1215 | 2 | 5 | Red maple : 22.6%<br>Yellow-poplar : 19.6%<br>White oak : 12.7% | Chestnut oak / black oak / scarlet oak : 13.3%<br>White oak / red oak / hickory : 25.1%<br>Yellow-poplar / white oak / northern red oak : 40.5% |
| CAR1215 | 24 | 86 | Eastern hemlock : 21.4%<br>Red maple : 17.6%<br>Scarlet oak : 10.1% | White oak / red oak / hickory : 19.5%<br>Chestnut oak / black oak / scarlet oak : 28.8%<br>Yellow-poplar / white oak / northern red oak : 30.9% |
| CAR1215 | 24 | 87 | Yellow-poplar : 19.5%<br>Red maple : 16.1% | Cherry / white ash / yellow-poplar : 10.0%<br>White oak / red oak / hickory : 18.9%<br>Chestnut oak / black oak / scarlet oak : 19.2%<br>Yellow-poplar / white oak / northern red oak : 35.4% |
| CAR1215 | 24 | 89 | Yellow-poplar : 17.6%<br>Sugar maple : 15.8%<br>Red maple : 10.7% | White oak / red oak / hickory : 29.4%<br>Yellow-poplar / white oak / northern red oak : 38.9% |
| CAR1215 | 24 | 90 | Chestnut oak : 33.0%<br>Red maple : 19.2% | Chestnut oak : 28.9%<br>Chestnut oak / black oak / scarlet oak : 60.6% |
| CAR1215 | 24 | 91 | Red maple : 22.6%<br>Yellow-poplar : 19.6%<br>White oak : 12.7% | Chestnut oak / black oak / scarlet oak : 16.6%<br>White oak / red oak / hickory : 26.8%<br>Yellow-poplar / white oak / northern red oak : 37.9% |
| ACR281 | 38 | 999 | Basswood : 10.8%<br>Red maple : 12.4%<br>Sugar maple : 36.0% | Hard maple / basswood : 19.6%<br>Sugar maple / beech / yellow birch : 79.4% |
| CAR1175 | 94 | 999 | Red spruce : 20.3%<br>Balsam fir : 14.2%<br>Eastern hemlock : 11.6%<br>Red maple : 10.8% | Red spruce : 29.0%<br>Sugar maple / beech / yellow birch : 50.6% |
| CAR1175 | 42 | 999 | Red spruce : 20.3%<br>Balsam fir : 14.2%<br>Eastern hemlock : 11.6%<br>Red maple : 10.8% | Red spruce : 28.4%<br>Sugar maple / beech / yellow birch : 51.3% |
| CAR993 | 55 | 221 | Douglas fir : 30.3%<br>Tanoak : 43.3% | Tanoak : 87.7% |
| CAR993 | 79 | 297 | Douglas fir : 30.3%<br>Tanoak : 43.3% | Tanoak : 91.9% |
| ACR378 | 79 | 297 | Tanoak : 59.6%<br>Douglas fir : 16.6% | Tanoak : 99.1% |
| ACR377 | 79 | 297 | Tanoak : 45.7%<br>Douglas fir : 33.4% | Tanoak : 90.7% |
| ACR260 | 25 | 999 | Douglas fir : 30.9%<br>Mountain hemlock : 18.8%<br>Grand fir : 13.3% | Douglas fir : 81.8% |
| ACR260 | 58 | 999 | Douglas fir : 30.9%<br>Mountain hemlock : 18.8%<br>Grand fir : 13.3% | Douglas fir : 85.8% |
| ACR371 | 41 | 999 | Eastern whitepine : 32.0%<br>Pin cherry : 16.0%<br>Sassafras : 11.6% | Eastern white pine / northern red oak / white ash : 37.1%<br>Sugar maple / beech / yellow birch : 62.9% |
| CAR1174 | 79 | 297 | Douglas fir : 52.9%<br>Tanoak : 13.2%<br>California black oak : 11.6% | California mixed conifer : 27.6%<br>Douglas fir : 54.0% |
| ACR417 | 42 | 167 | Black spruce : 13.1%<br>Northern white-cedar : 65.7% |  |
| ACR417 | 42 | 168 | Red maple : 35.4%<br>Red spruce : 11.0%<br>Yellow birch : 15.9% | Sugar maple / beech / yellow birch : 90.2% |
| ACR417 | 42 | 169 | Beech : 23.7%<br>Sugar maple : 21.8%<br>Yellow birch : 13.6% | Sugar maple / beech / yellow birch : 100.0% |
| ACR417 | 42 | 170 | Eastern hemlock : 12.8%<br>Eastern whitepine : 39.4% | Eastern white pine / eastern hemlock: 12.2%<br>Eastern white pine / northern red oak / white ash : 24.6%<br>Eastern white pine : 55.7% |
| ACR417 | 42 | 171 | Balsam fir : 10.1%<br>Paper birch : 33.7%<br>Quaking aspen : 18.2%<br>Red maple : 20.9% | Paper birch : 90.6% |
| ACR417 | 42 | 172 | Balsam fir : 27.2%<br>Red spruce : 23.3% | Red spruce : 11.9%<br>Balsam fir : 13.6%<br>Sugar maple / beech / yellow birch : 16.3%<br>Red spruce / balsam fir : 55.4% |
| ACR373 | 38 | 149 | Basswood : 11.7%<br>Red maple : 25.5%<br>Sugar maple : 30.2% | Sugar maple / beech / yellow birch : 92.7% |
| ACR282 | 79 | 297 | Douglas fir : 31.9%<br>Tanoak : 24.6%<br>Redwood : 20.6%<br>Red alder : 11.7% | Tanoak : 40.7%<br>Douglas fir : 49.1% |
| ACR282 | 55 | 221 | Douglas fir : 31.9%<br>Tanoak : 24.6%<br>Redwood : 20.6%<br>Red alder : 11.7% | Douglas fir : 33.9%<br>Tanoak : 58.5% |
| CAR1095 | 79 | 297 | Douglas fir : 41.0%<br>Ponderosa pine : 13.0%<br>California black oak : 13.0%<br>Southern scrub oak : 15.0% | California black oak : 14.8%<br>Douglas fir : 24.6%<br>California mixed conifer : 48.3% |
| ACR255 | 60 | 231 | Ponderosa pine : 29.6%<br>Douglas fir : 52.0% | Ponderosa pine : 17.4%<br>Douglas fir : 82.1% |
| ACR255 | 60 | 233 | Subalpine fir : 12.1%<br>Western larch : 20.9%<br>Lodgepole pine : 22.8%<br>Douglas fir : 29.8% | Engelmann spruce / subalpine fir: 13.3%<br>Lodgepole pine : 14.9%<br>Western larch : 17.9%<br>Douglas fir : 49.0% |
| ACR255 | 22 | 82 | Ponderosa pine : 44.8%<br>Douglas fir : 44.3% | Douglas fir : 43.4%<br>Ponderosa pine : 55.8% |
| CAR1103 | 55 | 221 | Douglas fir : 54.4%<br>Tanoak : 16.6% | Tanoak : 12.4%<br>California mixed conifer : 21.2%<br>Douglas fir : 56.1% |
| CAR1103 | 79 | 297 | Douglas fir : 54.4%<br>Tanoak : 16.6% | Tanoak : 12.8%<br>California mixed conifer : 17.2%<br>Douglas fir : 60.7% |
| ACR211 | 95 | 353 | Douglas fir : 15.0%<br>Ponderosa pine : 68.0% | Ponderosa pine : 99.0% |
| ACR211 | 95 | 350 | Douglas fir : 25.0%<br>Quaking aspen : 22.0%<br>Ponderosa pine : 13.0%<br>Engelmann spruce : 11.0%<br>White fir : 17.0% | Douglas fir : 90.3% |
| ACR324 | 286 | 286 | Alaska yellow-cedar : 14.1%<br>Sitka spruce : 13.9%<br>Western red cedar : 14.9%<br>Western hemlock : 40.2%<br>Mountain hemlock : 14.3% | Western hemlock : 90.9% |

## Footnotes

1 Assessment areas span the entire geography of their supersection. Despite the name, they represent not a geographic subset of areas but a subset of forest types that protocol developers deemed to have similar ecological and economic attributes.

## Notes

https://doi.org/10.5281/zenodo.4630712

https://doi.org/10.5281/zenodo.4630684

https://github.com/carbonplan/forest-offsets

https://github.com/carbonplan/forest-offsets-paper

### Competing Interest Statement

The authors have declared no competing interest.

